# Brain endothelial cells are exquisite sensors of age-related circulatory cues

**DOI:** 10.1101/617258

**Authors:** Michelle B. Chen, Hanadie Yousef, Andrew C. Yang, Davis Lee, Benoit Lehallier, Nicholas Schaum, Stephen R. Quake, Tony Wyss-Coray

## Abstract

Brain endothelial cells (BECs) are key elements of the blood-brain barrier (BBB), protecting the brain from pathogens and restricting access to circulatory factors. Recent studies have demonstrated that the circulatory environment can modulate brain aging, yet, the underlying processes remain largely unknown. Given the BBB’s intermediary position, we hypothesized that BECs sense, adapt to, and relay signals between the aging blood and brain. We sequenced single endothelial cells from the hippocampus—a brain region key to learning, memory, and neurogenesis— of healthy young and aged mice as well as post-exposure to inflammatory and age-related circulatory factors. We discovered that aged capillary BECs, compared with arterial and venous cells, exhibit the greatest transcriptional changes, upregulating innate immunity, antigen presentation, TGF-β signaling and oxidative stress response pathways. Remarkably, short-term infusions of aged plasma into young mice recapitulated key aspects of this aging transcriptome, while infusions of young plasma into aged mice reversed select aging signatures, essentially rejuvenating the BBB endothelium transcriptome. We identify candidate pathways mediating blood-borne brain rejuvenation by comparing age-upregulated genes with those modulated by plasma exposure. Together, these findings suggest that the transcriptional age of BECs is exquisitely sensitive to age-related circulatory cues and pinpoint the BBB itself as a promising therapeutic target to treat brain disease.

**Highlights:**

- Single-cell RNA sequencing of brain endothelial cells (BECs) reveals transcriptional segmentation into distinct arterial, capillary, and venous identities with age and experimental interventions
- Changes with age are heterogenous across vessel segments, with aged capillaries enriched in signatures of innate immunity, TGF-β and VEGF signaling, hypoxia and oxidative stress
- BECs sense and respond transcriptionally to diverse circulatory cues: inflammatory, pro-aging, or rejuvenating
- Aged plasma exposure recapitulates—and young plasma reverses—key transcriptomic signatures of normal BEC aging
- BEC response to aged and young plasma reveals cell non-autonomous mechanisms of blood-brain-barrier aging

## INTRODUCTION

Aging drives the deterioration of brain structure and function, increasing susceptibility to neurodegenerative disease and cognitive decline (Andrews-Hanna et al., 2007; Bishop et al., 2010; Mattson and Magnus, 2006). While the cell-intrinsic hallmarks of aging, from stem cell exhaustion to loss of proteostasis are established aspects of brain aging (López-Otín et al., 2013), recent studies have demonstrated cell non-autonomous mechanisms of brain aging via heterochronic parabiosis or blood plasma infusions (Wyss-Coray, 2016). Specifically, old plasma appears to impair and young plasma revitalizes cognitive function and hippocampal neurogenesis (Castellano et al., 2017; Katsimpardi et al., 2014; Khrimian et al., 2017; Villeda et al., 2011, 2014). Recently, infusion of aged plasma into young mice results in upregulation of vascular cell adhesion molecule 1 (VCAM1) in brain endothelial cells (BECs) and blocking via antibodies strongly reduces neuroinflammation and improves learning and memory function in aged mice (Yousef et al., 2019). Specific mouse and human proteins have recapitulated the effects of plasma, such as the pro-aging B2M and CCL11, and the rejuvenating TIMP2 (Castellano et al., 2017; Smith et al., 2015; Villeda et al., 2011). Nevertheless, while these studies show systemic effects on the brain, the exact mechanisms mediating these effects are largely unclear.

This is especially so considering that the brain is partitioned from the periphery via specialized vasculature—the blood-brain barrier (BBB) (Abbott et al., 2006; Broadwell, 1989; Daneman and Prat, 2014; Reese and Karnovsky, 2004). Relative to peripheral endothelium, the BBB exhibits limited permeability to macromolecules by employing unique tight junctions and low rates of transcytosis (Andreone et al., 2017; Ben-Zvi et al., 2014; Chow and Gu, 2015). These special properties are induced in development and maintained in adulthood by surrounding pericytes, smooth muscle cells, astrocytes, and neurons that form a functional ‘neurovascular unit’ (Armulik et al., 2010; Daneman et al., 2010). Dysfunction and breakdown of this unit have been implicated in age-related neurodegeneration and manifest in reduced cerebral blood flow, leakage of toxic factors, and a general inability to maintain an optimal environment for neuronal and stem cell function (Iadecola, 2013; Sweeney et al., 2018; Zhao et al., 2015; Zlokovic, 2008).

Though age-related BBB dysfunction has been probed with a diverse toolkit of tracers, the transcriptional heterogeneity of the BBB and vessel segment-specific responses to the parenchymal or systemic environment has been largely unexplored (Bien-Ly et al., 2015; Marques et al., 2013; Montagne et al., 2015; Mooradian, 1988; Vanlandewijck et al., 2018). Here, we study normal brain endothelial aging—and its response to inflammatory and age-related circulatory cues—by profiling hippocampal BECs using single-cell RNA sequencing. We characterize significant transcriptional changes across arterial, capillary, and venous cells, discovering a surprising malleability to age-related plasma factors, and a heterogenous distribution of age-related receptors and signaling pathways across vessel segments. This suggests the BBB endothelium is positioned to and capable of mediating reversible, non-autonomous mechanisms of brain aging.

## RESULTS

### Brain endothelial cells exhibit segmental identities

We rapidly isolated and pooled CD31+CD45-Cd11b-BECs from mouse hippocampi and analyzed their transcriptomes using single-cell RNA sequencing as previously described (**Figure 1A, Figure S1A-C)**(Yousef et al., 2019). Cells passing QC had at least 50,000 reads, with a median of ∼700,000 reads and ∼1,800 expressed genes per cell (**Figure SI 1D-E**). All cells expressed at least one pan-BBB/endothelial marker at the mRNA level (*Cldn5, Cdh5, Pecam1, Ocln, Flt1, Esam*). Few cells exhibited both high mitochondrial and ribosomal gene counts, typical features of poor cell quality or health during the isolation and collection phase (Butler et al., 2018) (Figure SI 1F).

**Fig 1.**
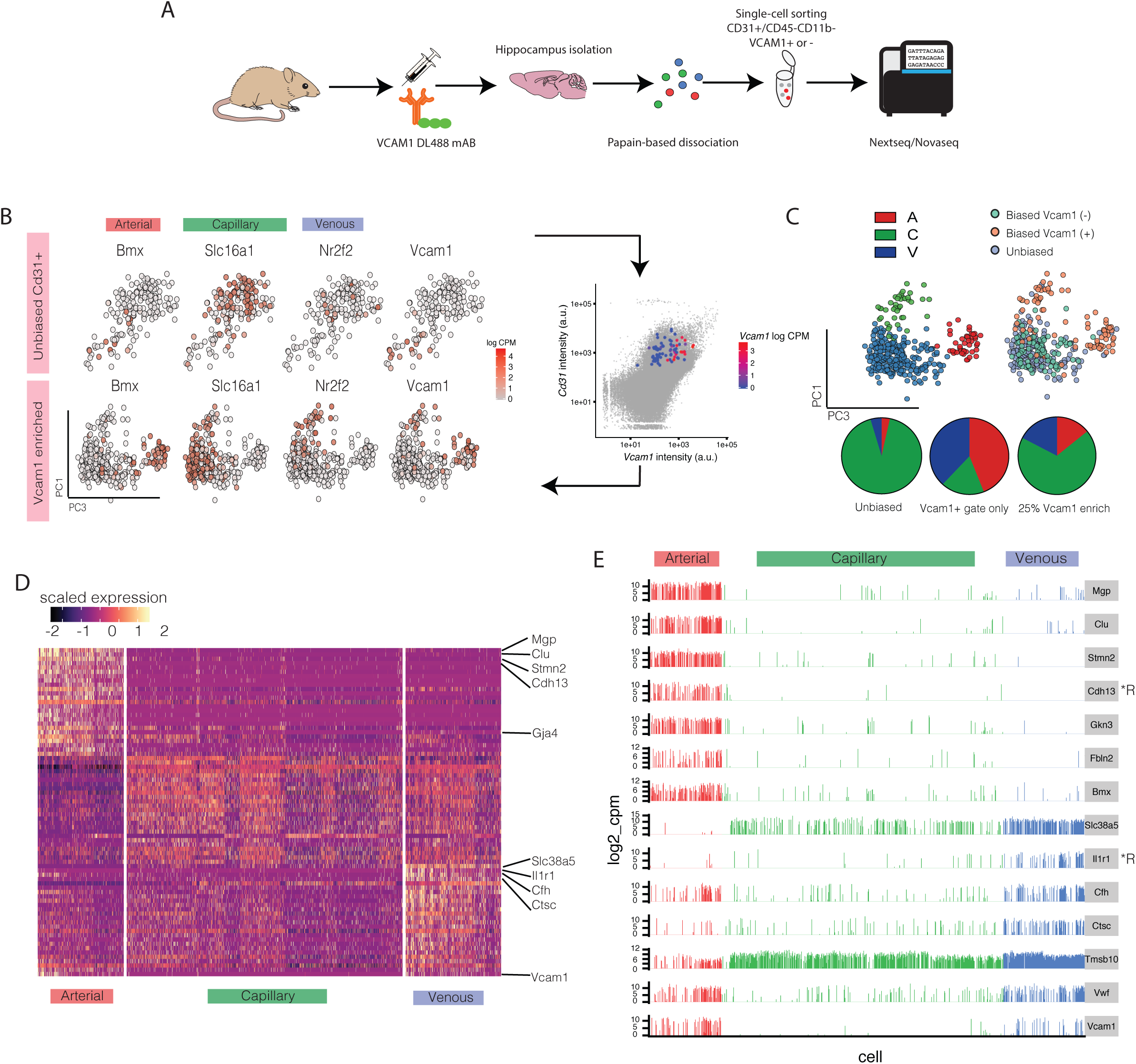
Brain endothelial cells segment into arterial, capillary, and venous identities. a. Schematic of experimental protocol for single-cell analyses of BEC transcriptome. b. (Top panel) tSNE of a subset (179) of 3 m.o. BECs collected in an unbiased manner (CD31+) and non-discrete expression pattern (*logcpm*) of key A-C-V genes including *Bmx* (arterial), *Slc16a1* (capillary), *Nr2f2* (venous), and *Vcam1* (arterial & venous). Note the low number of *Vcam1* + cells and the absence of a clear venous EC population. (Bottom panel) Addition of a subset of *Vcam1*+ cells into PCA analysis (62, as collected by FACs enrichment) significantly improves A-C-V identification. c. Calling of arterial, venous and capillary populations after the addition of *Vcam1*+ cells. Pie charts of the proportion of ACV cells in unbiased CD31 sorts, in VCAM1+/CD31+ sorts, and unbiased with 25% of VCAM1+ cell enrichment (final condition). d. Heatmap of the top 25 most enriched genes per A-C-V population (which were identified via unbiased whole transcriptome clustering) in young BECs. e. Barplots of the expression level of top genes which may act as novel markers for A and V identities. Genes encoding for cell surface receptors are indicated by *R. Expression levels in A-C-V segments are validated for the full set of 981 3 m.o. BECs (over 4 biological replicates).

We first characterized the range of distinct cell populations within heterogenous hippocampal BECs from young (3 month-old) mice via transcriptome clustering of the top 2,500 over-dispersed genes. Specifically, we searched for BEC populations defining segmental identities of arterioles, capillaries, and venules, as previously shown at a single-cell level (Vanlandewijck et al., 2018). Principal component analyses did not yield clear segmental or other phenotypic signatures, with venous (V) (*Slc38a5, Nr2f2*) and capillary (C) (*Slc16a1*) markers showing a generally diffuse distribution along the first 10 PCs, and arterial (A) (*Bmx, Efnb2*) markers being slightly more biased **(Figure 1B, top panel)**. Upon further inspection, we found that FACS sorting via CD31+/CD45-alone yielded low numbers of arterial and venous cells (<6% per population), which were defined by a non-zero expression of at least 2/3 classical A and V gene markers (Arterial: *Bmx, Efnb2, Vegfc*, Venous: *Nr2f2, Slc38a5, Vwf*) **(Figure S2A)**. Vascular cell adhesion molecule (*Vcam1)*, a cell surface receptor that facilities endothelial-immune cell interactions, has previously been shown to be highly enriched in arterial and venous cells and we established a method to isolate and enrich primary venous and arterial BECs using this marker (Vanlandewijck et al., 2018; Yousef et al., 2019). Taking advantage of the surface expression of VCAM1, we infused a fluorescently labeled anti-VCAM1 mAb retro-orbitally prior to mouse perfusion and tissue dissection which allowed us to enrich VCAM1+BECs using FACS. Addition of VCAM1+ sorted cells to the original dataset (∼25% of all cells) resulted in a more biased (yet still continuous) distribution of the expression of known A-C-V markers, and an increase in A and V cell identities (**Figure 1B-C**). VCAM1 protein levels were highly correlated with mRNA content, and, nearly all *Vcam1* mRNA+ cells were co-positive for and highly correlated to either A or V markers, and largely absent in capillaries (**Figure SI 2B**). Not all arterial and venous cells defined were *Vcam1*+, suggesting that *Vcam1* is only expressed in a subset of arterial and venous cells, Indeed, differential expression tests between Vcam1+/-arterial or venous populations show a basally more transcriptionally activated subset of BECs (**Figure SI 2C**) that are confined within A and V populations. Clarity of A-C-V populations was improved due to the increase in number of A and V cells (**Figure SI 2D**), which allows for previously small populations of arterial and venous cells, some of which expressed gene signatures more similar to capillaries on the zonated A-C-V gene expression axis, to emerge (Vanlandewijck et al., 2018).

Furthermore, by finding genes which are most enriched in arterial and venous clusters, we were able to identify potential new segmental markers for BECs **(Figure 1D)**. Venous cells exhibited more shared genes with capillaries, than arterial cells. Arterial cells were enriched in *Mgp, Clu, Stmn2, Cdh13*, while venous cells were enriched in *Il1r1, Cfh, Ctsc and Tmsb10*. In fact, in contrast to classical venous (*Nr2f2*) and arterial (*Efnb2*) markers, these new markers were expressed across a significantly larger number of cells in their respective segment populations. In addition, these genes are not restricted to expression in *Vcam1+* subpopulations, making them more suitable markers for pan-arterial/venous cell identification (**Figure SI 2C**). Of note, gene products of *Cdh13* (Cadherin-13) and *Il1r1* (Interleukin 1 Receptor Type 1) are known to be expressed on the cell surface and confirmed to be enriched in the hippocampus, making them potential candidates for FACS enrichment of arterial or venous cells **(Figure 1E**).

### Systemic LPS administration activates common transcriptional programs across segment identities

To understand whether BECs can act as sensors of organismal-level perturbations, we administered LPS systemically in young mice to induce an acute inflammatory response. LPS serves as an acute perturbation, where dramatic organismal-wide changes are expected, and thus facilitates a preliminary study of BEC response to systemic cues. Out of 10,955 expressed genes across all BECs, a total of 1,610 differentially expressed genes (DEGs) were identified (FDR<0.05 threshold) between LPS-treated and untreated mice, with 865 DEGs in capillaries, 881 in venous, and 956 in arterial identities. 357/1610 (22%) DEGs are shared between all three segments, while some are unique to one or two segment identities. Fairly even numbers of up- and down-regulated genes are observed with LPS, for all three segments **(Figure 2A).** Furthermore, LPS stimulation did not seem to change the native compositions of A-C-V identities (**Figure SI 3A**). GO pathway analyses of both up- and down-regulated DEGs reveal largely common pathways between vessel segments, including the upregulation of interleukin and interferon signaling, cytoskeletal remodeling, cell-matrix adhesion, and TGF-β signaling pathways, as well as the downregulation of EC proliferation, lipid and lipoprotein metabolism, and adherens junctions maintenance **(Figure 2B)**. LPS induced large fold changes in expression levels of DEGs, with many genes exhibiting on-off responses such as the innate immunity genes *Lcn2, Icam1, Cebpd, Irf7, Litaf, Ifit3* **(Figure 2C)**. *Lcn2* (Lipocalin2), a neutrophil-associated lipocalin that plays roles in innate immunity, was the most highly upregulated gene following LPS treatment in all A-C-V segments, while *Cd14*, a receptor for LPS was significantly upregulated in venous cells.

**Fig 2.**
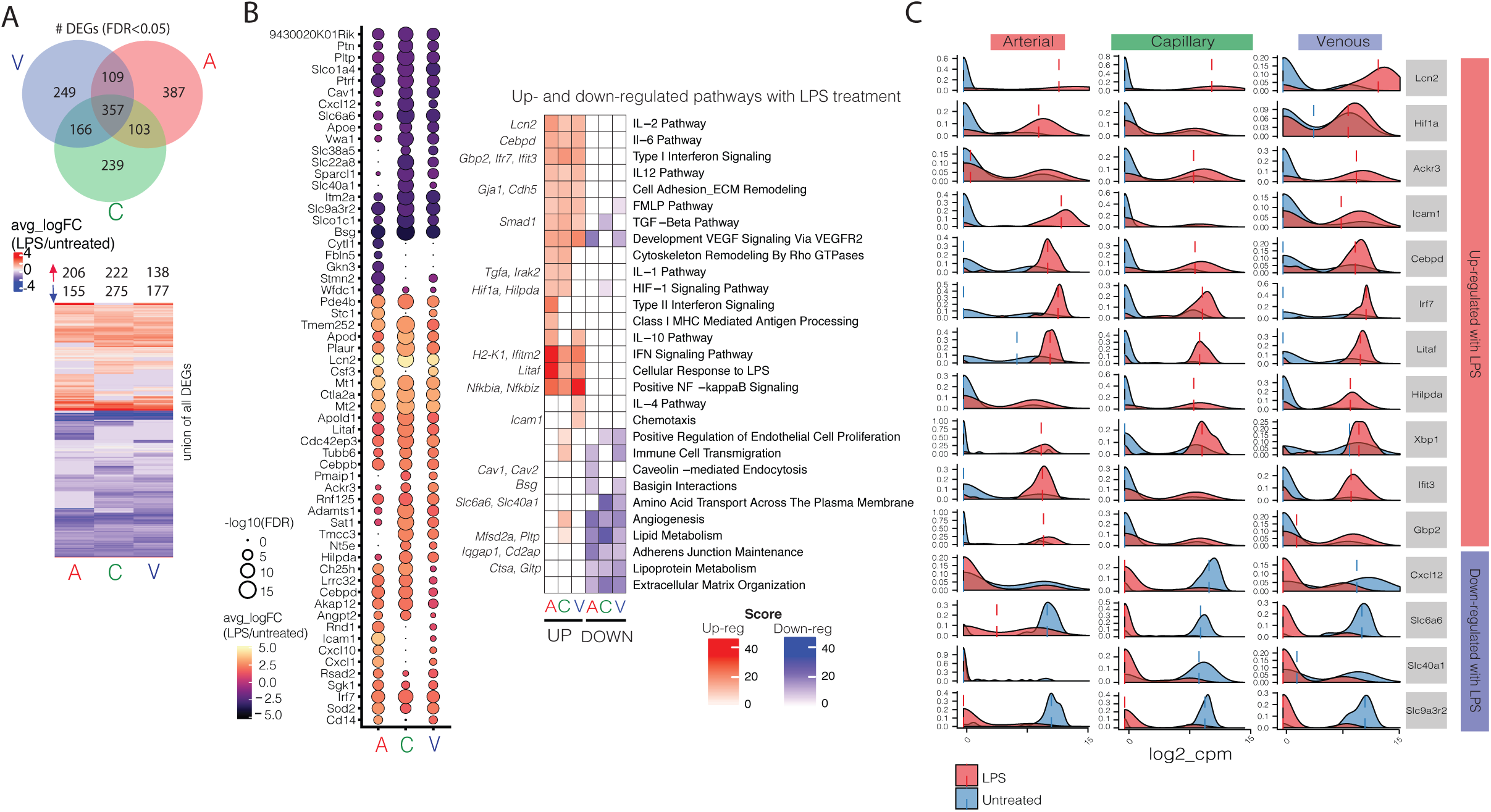
Systemic LPS administration activates common transcriptional programs across segment identities. a. Venn diagram showing the number of DEGs (FDR<0.05) shared between each vessel segment. Heatmap showing the distribution of up- and down-regulated genes per vessel segment. Dotted heatmap of top 60 DEGs ranked by avg_logFC*-log10(FDR). Color indicates the average log fold change of LPS/untreated, while the dot size represents degree of statistical significance. Only genes with FDR<0.05 for at least one vessel segment is listed, and hierarchical clustering is performed (dot size = 0 indicates FDR>0.05). b. GO enrichment analysis of the list of DEGs (FDR<0.05) up- and down-regulated in LPS treated A, C and V cells. Left hand side (red) indicates pathways that are over-represented by DEGs upregulated with LPS, right hand side (blue) indicates pathways over-represented by DEGs down-regulated with LPS treatment. Exemplary genes contributing to pathway enrichment in the upregulated DEG set is listed on the side. c. Density plots of key genes from (a) showing the single cell distributions of expression levels in A, C and V segments. Dotted lines indicate median of the LPS or untreated sample distributions. All comparisons shown between LPS and untreated are significant (p<0.05).

### Acquisition of aging BEC transcriptomic signatures is distinct across vessel segments

Aging results in prominent changes in brain function and the hippocampus appears particularly vulnerable, showing the first signs of degeneration in Alzheimer’s disease (Wyss-Coray, 2016). Because BECs are responsible for nutrient transport into the brain and communication between peripheral immune cells and the CNS, understanding how they age is crucial to understanding brain aging and neurodegeneration. We sequenced CD31+/CD45-/CD11b-BECs from the hippocampi of young (3 month-old) (981 cells) and aged (19 month-old) (1053 cells) disease-free mice. Approximately 20% of BECs were enriched for VCAM1 expression by FACS to increase the collection of arterial and venous BECs. Unbiased transcriptome clustering of all young and aged cells combined revealed 3 continuous subpopulations with transcriptional signatures of A-C-V identities, illustrated by the gradual zonation of *Gja4, Bmx, Slc16a1, Slc38a5, Nr2f2, and Vcam1* **(Figure 3A-B, Figure SI 4)**. Aged and young BECs did not appear to show clear distinguishing signatures within the first 10 PCs (**Figure SI 5**), indicating that age does not obviously alter segmental identity.

**Fig 3.**
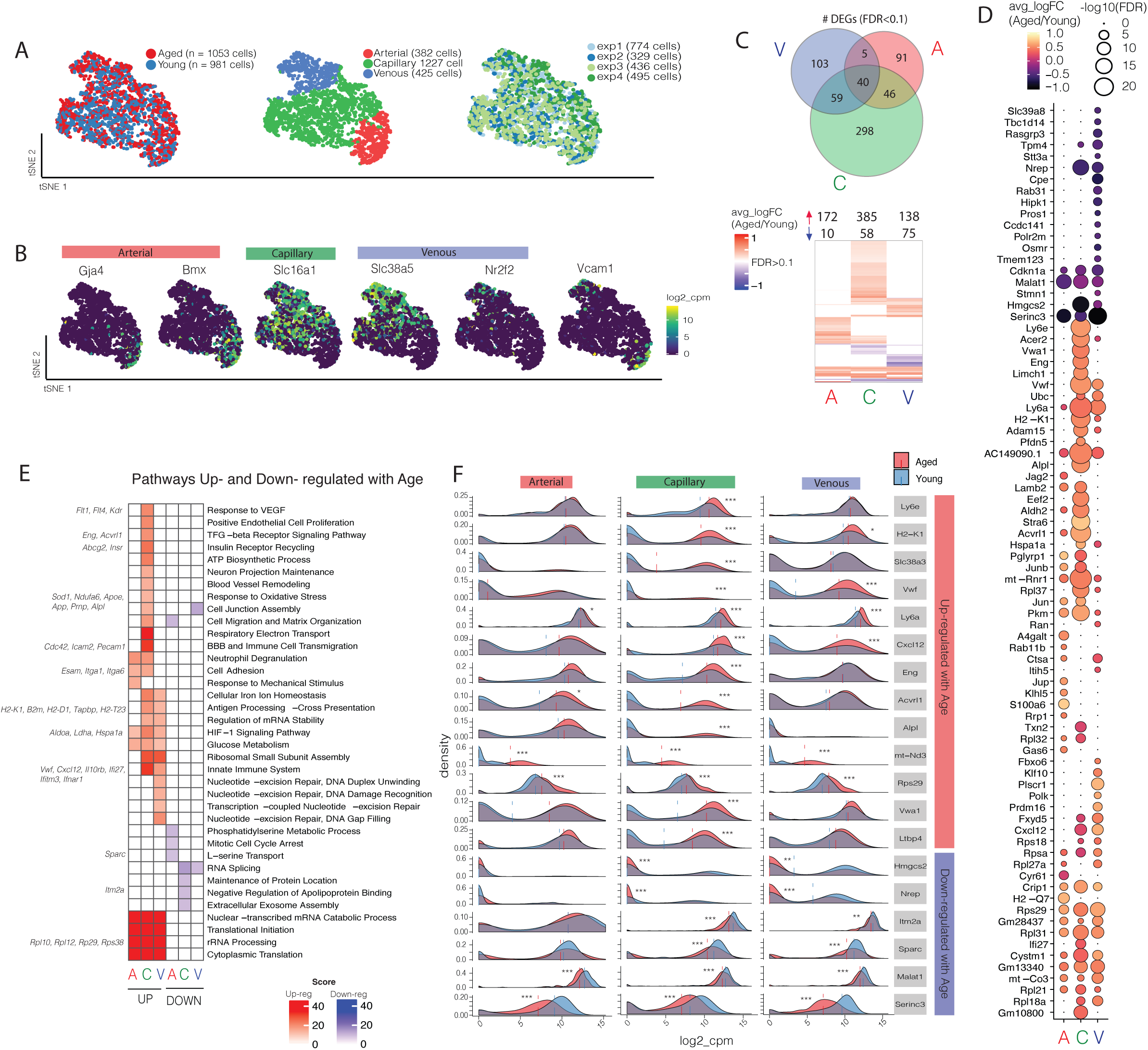
Healthy aging of BECs results in transcriptome changes that are distinct across segment identities. a. tSNE after aligning healthy aged and young datasets via Canonical Correlation Analysis (CCA), using the top 9 correlation components. Aged and young cells show comparable distribution of A, C, and V identities along the A-C-V axis. Note that segmental identity largely drives cluster formation, rather than age group. b. Distribution of key A-C-V marker genes in tSNE-space. c. Venn diagram showing the numbers of DEGs (FDR<0.1) shared between different vessel segments. Heatmap of the union of all DEGs up- and down-regulated in aged A, C, or V cells, illustrating the degree of overlap of DEGs between each segment. d. Dotted heatmap of top 80 DEGs ranked by avg_logFC*-log10(FDR) (a subset of genes in (c). Color indicates the average log fold change of Aged/Young, while the dot size represents degree of statistical significance. Any gene with FDR<0.1 for at least one vessel segment is listed, and hierarchical clustering is performed (dot size = 0 indicates FDR>0.1). d. GO analysis of all DEGs (FDR<0.1) up- and down-regulated in aged A, C and V cells. Left hand side (red) indicates pathways over-represented by genes upregulated with aging, right hand side (blue) indicates pathways over-represented by genes down-regulated with aging. Exemplary genes contributing to pathway enrichment in the upregulated DEG set is listed on the side. e. Density plots of key genes from (d) showing the single cell distributions of expression levels in A, C and V segments. Dotted lines indicate median of the young or aged distribution. *p<0.1, **p<0.01, ***p<0.001.

Comparisons of aged and young BECs within A-C-V populations results in a total of 642 unique DEGs (FDR<0.1, capillary: 443, venous: 207, arterial 182 DEGs). Interestingly, the degree of overlap in DEGs within vessel segments was much less compared to LPS treatment **(Figure 2)**, with only 40/642 (6%) DEGs shared between all three vessel segments. The majority of DEGs were found to be increased (86%) rather than decreased, suggesting a general upregulation of transcriptional programs (**Figure 3C**). Capillary cells exhibited higher numbers of DEGs than arterial or venous cells, with 298/443 (67%) of their DEGs being unique to the segment only **(Figure 3C-D)**. Interestingly, aged cells exhibited a slightly higher number of expressed genes (mean=1,801 compared to 1,474), while capillaries expressed ∼25% fewer genes than arterial and venous cells (mean=1,465 compared to A: 1,899 and V: 1,928) (**Figure SI 6A-B**).

Aged capillaries reveals strong upregulation in genes including stem-cell antigen 1 and 2 (*Ly6e, Ly6a)*, innate immunity (*Vwf, Cxcl12, Dusp3, Ifi27, Ifnar1, Il10rb*), antigen-processing (*B2m, H2-K1, H2-D1, Tapbp, H2-T23*), VEGF-signaling (*Kdr, Flt1, Flt4)*, matrix assembly (*Vim, Vwa1, Spock2)*, cell adhesion (*Itga1, Itga6, Esam*), TGF-β signaling (*Eng, Acvrl1, Ltbp4*), hypoxia response (*Ldha, Pkm, Aldoa, Nos3*), and oxidative stress (*Sod1, Apoe, App, Prnp, Alpl*) **(Figure 3E-F)**. We also find a strong and consistent upregulation of genes encoding ribosomal subunits across all segments (e.g. *Rpl37, Rpl31, Rpl21, Rpl35, Rplp2, Rpl37a, Rps20, Rps27a*) (**Figure 3E**). Changes in gene expression levels between aged and young BECs are subtler than those after LPS treatment. A comparison of DEGs in disease-free aging and with LPS treatment reveals few commonly shared DEGs, however several involved in innate immunity were commonly upregulated, including *B2m, H2-K1, H2-D1*, as well as some involved in ribosomal biogenesis and rRNA processing (*Rpl23, Rps12, Rps27, Rpl10, Rpsa*) (**Figure SI 3B**).

To ensure that the DEGs were not a consequence of differing cell numbers between tested groups or biological noise, we performed two sets of stringent tests. A permutation test was conducted on all DEGs (FDR<0.1) to ensure that the true average log fold change of each DEG fell beyond the 95th percentile of a randomly shuffled distribution (**Figure SI 6C**). In addition, DEGs were calculated within each of the four biological replicates (one biological replicate consisting of 4 pooled mice hippocampi), and only those found to be differentially expressed in 3 out of 4 replicates passed the criteria. Altogether, we find that each vessel segment ages differentially, and that aged capillaries exhibit the greatest degree of change, upregulating signatures such as innate immunity, antigen processing, TGF-β signaling, and oxidative stress response.

### Systemic injection of young mice with aged plasma recapitulates key signatures of aging in BECs

An aged circulatory environment, including changes in plasma or CSF proteomes, can promote brain dysfunction (Silva-Vargas et al., 2016; Villeda et al., 2011). However, the cellular and molecular mechanisms involved in relaying circulatory signals into the brain are unclear. We hypothesized that BECs play an intermediary role in sensing and responding to an aged circulatory proteome. Thus, we measured the transcriptional response of young BECs to soluble factors in the plasma of aged mice. We injected young mice with pooled plasma from aged mice (AMP) or PBS (150 ul per injection) retro-orbitally, twice-daily for 4 days (Yousef et al., 2019) and collected CD31+/CD45-as well as CD31+/CD45-/VCAM1+ cells from the hippocampus (**Figure 4A**). Single cell RNA sequencing and dimensionality reduction of AMP (n=333 cells) and PBS (n=206) treated young BECs revealed the same arteriovenous zonation found in normal aged mice, with segmental identity, rather than plasma treatment, being the main driver of heterogeneity (within the first 15 PCs) **(Figure 4B-C, Figure SI 7A).** Interestingly, BECs again respond differentially to plasma treatment depending on the vessel segment identity, with capillaries exhibiting a strikingly larger number of DEGs compared to arterial and venous cells, even when they are downsampled to match sample powers in other groups. Out of 12042 detected genes, 829 genes were found to be differentially regulated in capillaries (FDR<0.1) and most are up-regulated (692 – 83% of DEGs). Importantly, only a small subset (<10%) of these DEGs were also found differentially perturbed by injecting aged-matched young plasma into young mice, indicating that most of the effects of AMP in young mice are specific to the age of the plasma (SI Table 1). Thus, capillaries are highly responsive to factors in the exogenous aged plasma, inducing activation of existing or new transcriptional programs **(Figure 4D-E, Figure SI 7D**).

**Fig 4.**
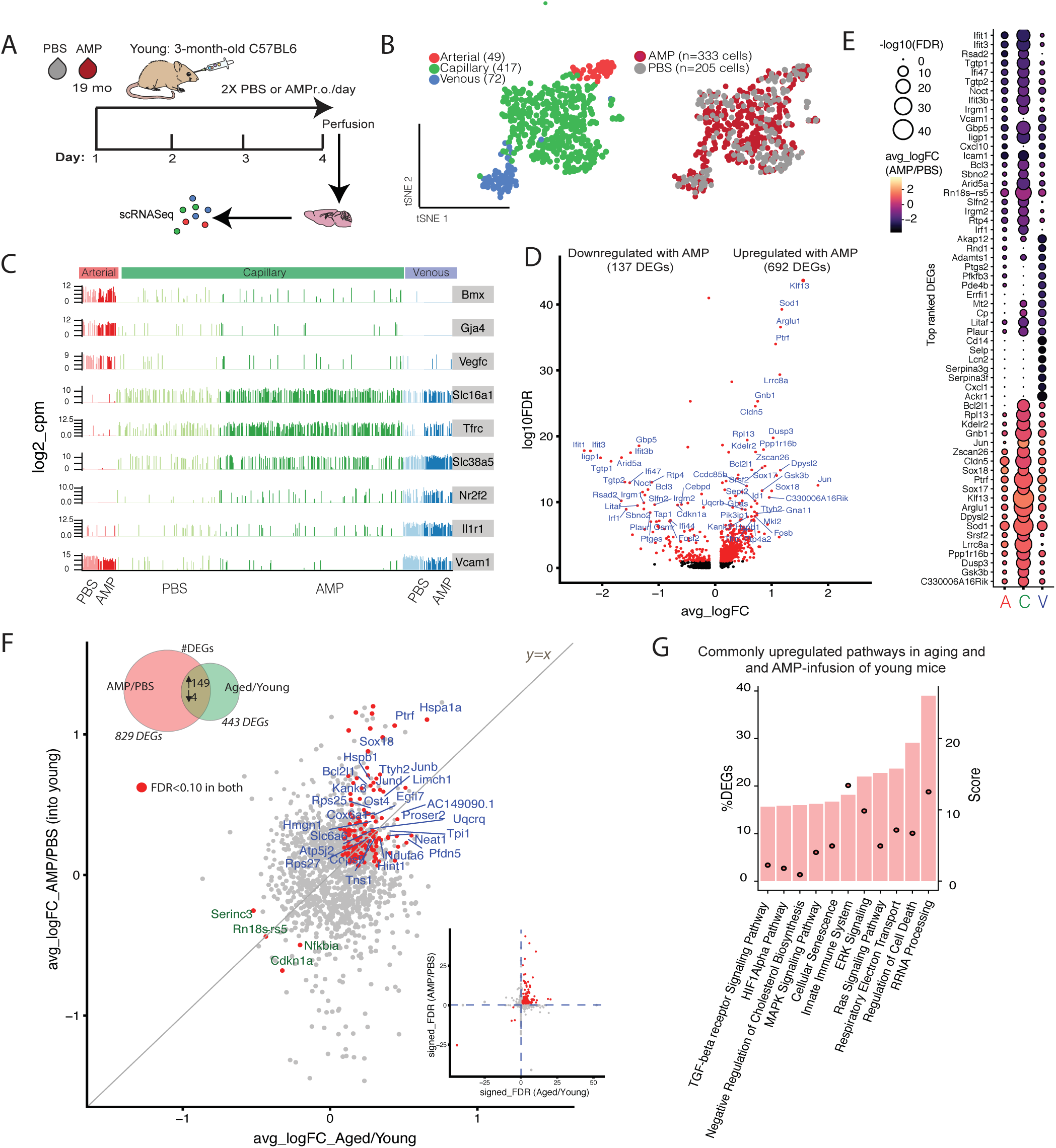
BECs sense cues in the circulatory milieu - aged plasma recapitulates signatures of healthy aging. a. Schematic of the AMP acute infusion into young mice paradigm. b. tSNE after aligning AMP and PBS treated datasets via Canonical Correlation Analysis (CCA), using the top 9 correlation components (CCs). AMP and PBS treated cells show comparable distribution of A, C, and V identities along the A-C-V axis, suggesting plasma infusions do not significantly alter native segmental identities. c. Barplot of expression level of canonical A-C-V marker genes in all AMP and PBS treated BECs, with segmental identity defined by unbiased clustering in (b). d. Volcano plot depicting up- and down-regulated genes with AMP treatment in capillaries only (compared to PBS control). Genes marked in red are significant (FDR<0.1). FDR values are calculated only with genes showing an avg_logFC>0.1. Genes labeled red are FDR<0.1. e. Dotted heatmap of top 60 DEGs ranked by avg_logFC*-log10(FDR) (a subset of genes in (c). Color indicates the average log fold change of AMP/PBS, while the dot size represents degree of statistical significance. Any gene with FDR<0.1 for at least one vessel segment is listed, and hierarchical clustering is performed (dot size = 0 indicates FDR>0.1). f. Scatterplot of genes and their log fold change in both healthy Aged/Young and AMP/PBS treatment in capillaries. The 153 genes that are commonly up- (blue) or downregulated (green) in both groups (and satisfy FDR<0.1 in both), are labeled. These genes are more likely ones that are modulated by aged plasma factors in a normal aged milieu, rather than ones specifically upregulated by plasma treatment, and unrelated to normal aging. Inset shows the same genes (red) on a plot of the signed-FDR value (sign of logFC*-log10(FDR)). g. Top pathways represented by the genes which are both upregulated in healthy aging and AMP treatment (149 intersecting DEGs).

Excitingly, we discovered a significant overlap between transcriptional changes in BECs as a result of normal aging and exposure to AMP. This overlap was most striking in capillaries and less pronounced in venous and arterial cells (**Figure SI 7B-C**). Out of 443 DEGs in aging and 829 DEGs in AMP treatment, 149 (up-regulated) and 4 (down-regulated) transcripts were found to be shared, and these intersecting DEGs comprised ∼34% and 18% of total DEGs in each comparison, respectively **(Figure 4F)**. This overlap is significant as the intersecting number is above the 99^th^ percentile of the distribution of intersects if genes were randomly chosen from each group. Surprisingly, nearly all of the intersecting genes are expressed at higher levels in AMP treated cells compared to normal aging **(Figure SI 7E-F)** suggesting that factors in AMP are powerful inducers of key aspects of BEC aging. Indeed, pathway analysis of the 149 commonly up-regulated transcripts pointed to similar pathways enriched in normal aging, including innate immunity (*Dusp3, Ifi27, Ifnar1, Il10rb, Vim, H2-T23, Icam2, Calm1, Myo10, Anxa2, Canx*), cellular senescence (*Uba52, Sod1, Rbx1, Elob, Fkbp4*), TGF-β signaling (*Nedd8, Bmpr2, Id1, Pdgfb*), hypoxia and stress (*Hspa1a, Hspb1*), and ribosomal processing (*Rpl10, Rpl10a, Rpl13, Rpl18a, Rpl26, Rpl28*) (**Figure 4G**).

### Systemic injections of aged mice with young plasma reverses key transcriptional changes of aging in BECs

The plasma of young mice can exert rejuvenating effects on the brains of aged mice after intravenous delivery, resulting in increased neurogenesis, memory and learning, dendritic spine density, and decreased neuro-inflammation and microglial activity (Wyss-Coray, 2016). To test if aged BECs are similarly responsive to acute injections of young mouse plasma (YMP), we injected aged mice with pooled YMP or PBS, isolated hippocampal BECs, and sequenced RNA from individual cells as described above (**Figure 5A**). Dimensionality reduction of all YMP (n=256) and PBS control treated cells (n=121) resulted in distinct ACV populations, with no obvious separation between treatment conditions (PCs 1 to 15) (**Figure 5 B-C**). Again, capillaries responded most significantly to plasma injections but, unlike BECs exposed to AMP, the great majority of transcripts were downregulated with YMP infusion (206/257 DEGs – 80% downregulated) (**Figure 5D**). Prominently down-regulated pathways include antigen processing and presentation via MHC Class 1 (*H2-D1, H2-Q6, H2-Q7, H2-T22, H2-T23, B2m, H2-K1, Tapbp*), innate immune response and cytokine (interferon) signaling (*Icam2, Ifi27, Ifitm3, Ifih1, Ifit3, Vwf*,) metabolic processes, and ribosomal biogenesis and rRNA processing (*Rpl13, Rpl38, Rpl41, Rps27, Rps27a, Rps29, Rps8*) (**Figure SI 8A**).

**Fig 5.**
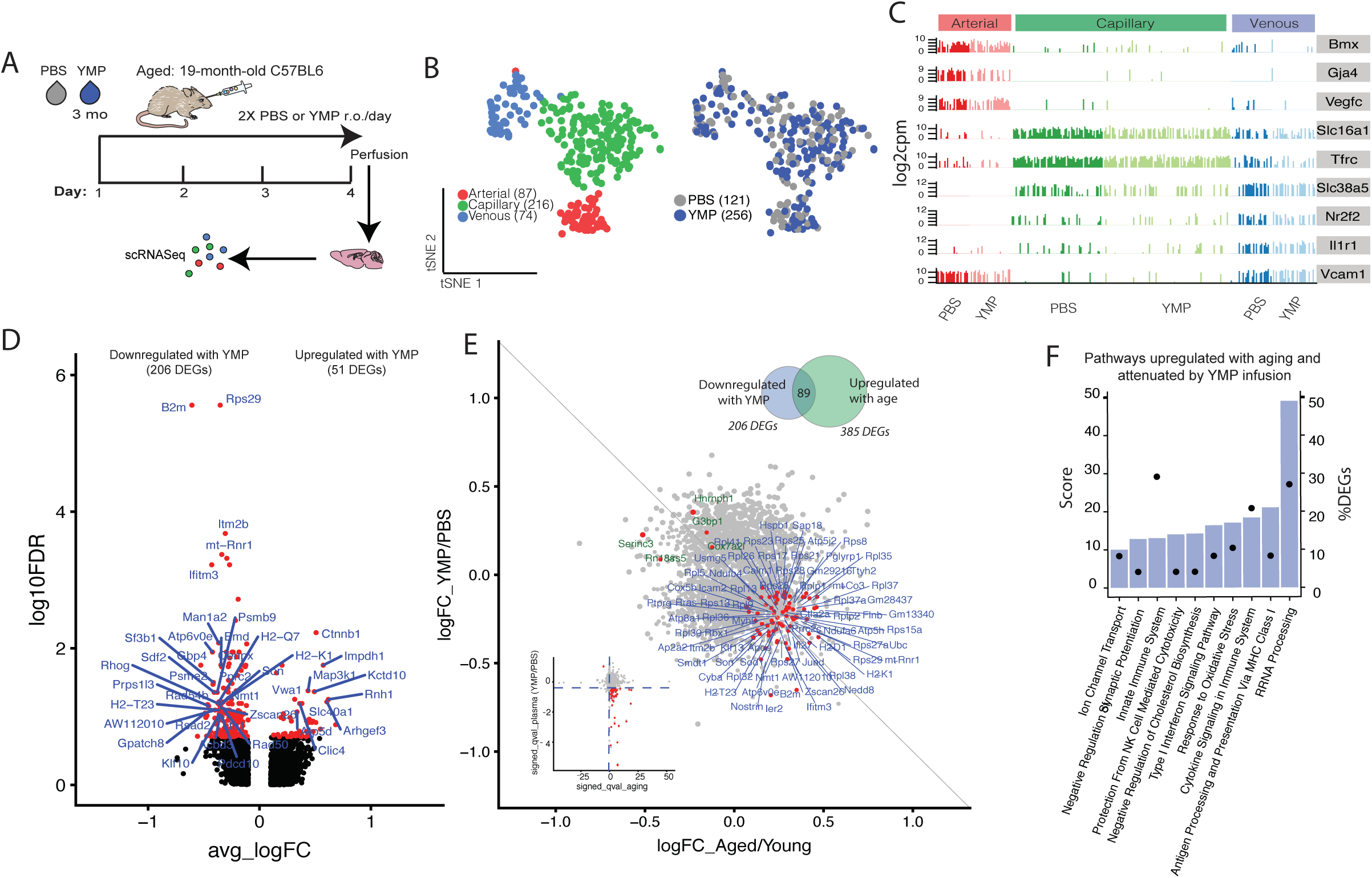
YMP plasma infusion reverses select signatures of normal BEC aging. a. Schematic of the YMP acute infusion into aged mice paradigm. b. tSNE after aligning YMP and PBS treated datasets via Canonical Correlation Analysis (CCA), using the top 9 correlation components (CCs). YMP and PBS treated cells show comparable distribution of A, C, and V identities along the A-C-V axis, suggesting plasma infusions do not significantly alter native segmental identities. c. Barplot of expression level of canonical A-C-V marker genes in all YMP and PBS treated BECs, with segmental identity defined by unbiased clustering in (b). d. Volcano plot of DEGs up- and down-regulated with YMP treatment (vs PBS control). FDR values are calculated only with genes showing an avg_logFC>0.1. Genes labeled red are FDR<0.1. e. Scatterplot of genes and their log fold change in both healthy Aged/Young and YMP/PBS treatment in capillaries. The 89 genes that are upregulated with age (Aged/Young), but decreased with YMP (YMP/PBS) and vice-versa (and satisfy FDR<0.1 in aging and FDR<0.1 in YMP), are labeled. It is likely that these genes upregulated in normal aging are able to be modulated and/or reversed with exposure to YMP. Inset shows the same genes (red) on a plot of the signed-FDR value (sign of logFC*-log10(FDR)). f. Top pathways represented by the genes which are upregulated in healthy aging and downregulated with YMP treatment (89 intersecting DEGs).

These findings suggest that YMP infusions are capable of reversing certain BEC aging signatures. Indeed, in capillaries, 89 DEGs increase with normal aging and decrease following YMP infusion, which comprises 12% and 31% of DEGs in normal aging and YMP infusions, respectively (**Figure 5E**). Strikingly, these 89 genes are enriched in key aging signature pathways (**Figure 3E**) including ribosomal biogenesis/rRNA processing (*Rpl13, Rpl31, Rpl36, Rpl38, Rpl41, Rps13, Rps21, Rps27, Rps28, Rps8*), immune system and cytokine signaling (*Vwf, Ifi27, Ifitm3, Ifitm2, Ifit3*), antigen processing and presentation (*B2m, H2-K1, H2-D1, H2-T23, Psmb9, Psmc2*), and response to oxidative stress (*Ndufb4, Apoe, Sod1, Nostrin*) (**Figure 5D**).

### Young plasma reverses select transcriptional changes of aging induced by AMP

To determine whether young plasma factors could specifically reverse transcriptional changes in BECs induced by aged circulatory factors, we compared the 149 shared DEGs between Aged/Young and AMP/PBS (**Figure 4F**) and 89 shared DEGs between Aged/Young and YMP/PBS (**Figure 5E**) using GeneAnalytics software to identify commonalities in pathways and directionality (**Figure 6A-B, Figure SI 9**). Pathways represented in both datasets include TGF-β signaling, cellular senescence, respiratory electron transport, innate immunity, interferon signaling, cholesterol biosynthesis, response to oxidative stress, and rRNA processing. It is important to note that genes enriched in each common pathway do not entirely intersect, suggesting that upregulation and then attenuation of pathways may not necessarily involve the same full set of genes. 42 DEGs lie in the “triple-intersect”, representing various pathways such as immune response signaling, with antigen processing (*H2-T23*), immune cell adhesion (*Icam2*), and interferon (*Ifi27*) amongst the top genes upregulated by aging and AMP and downregulated by YMP (**Figure 6C**). Strikingly, oxidative stress response was strongly enriched in aging and AMP-treated BECs and reduced by YMP, with *Apoe, Sod1, Ndufa6, Nostrin*, and *Selenow* being differentially regulated in all three datasets.

**Fig 6.**
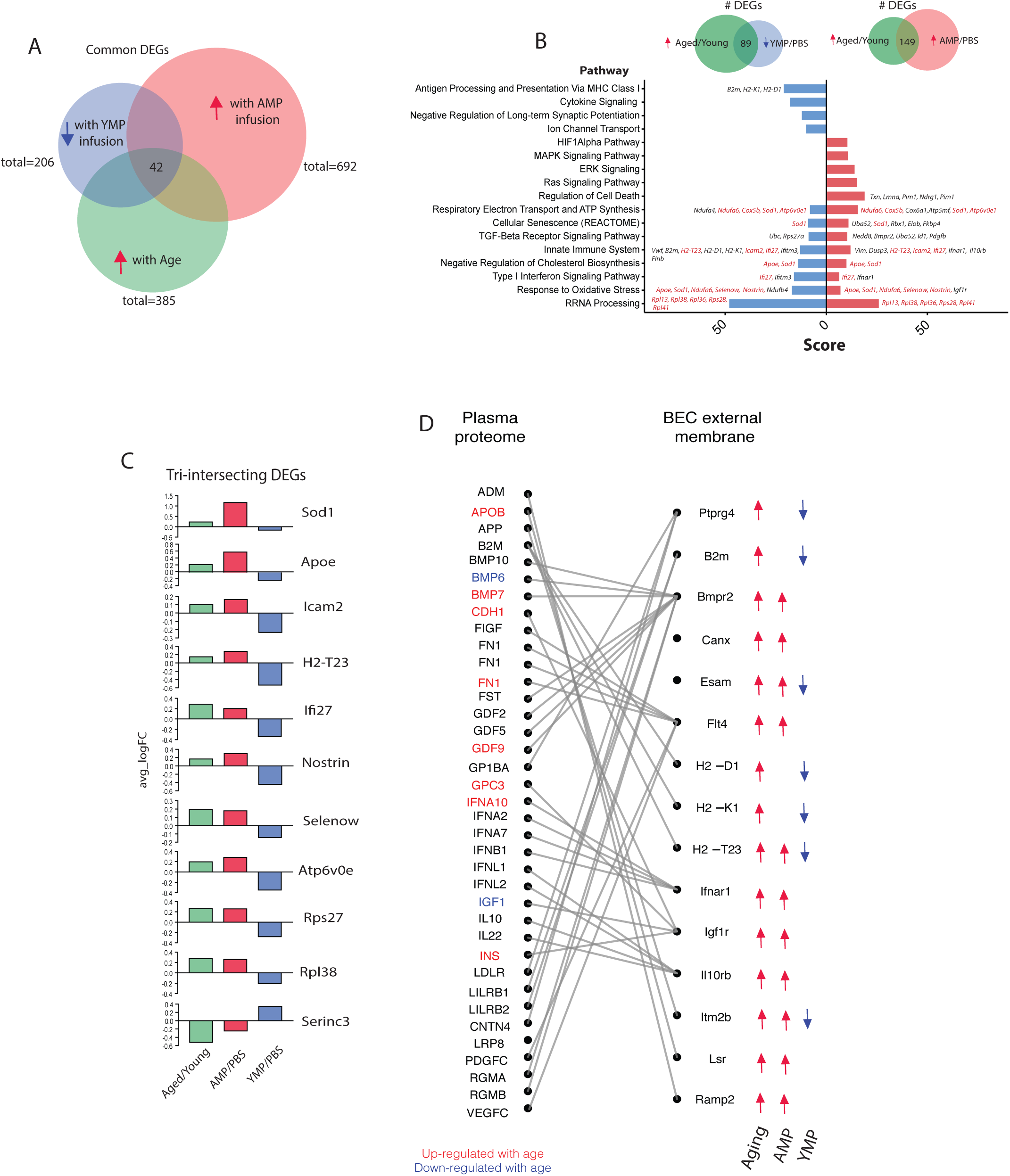
Young plasma administrations can rejuvenate BECs aging signatures that are induced by aged plasma. a. Venn diagram depicting the number of DEGs shared between each treatment condition (Aged/Young, AMP/PBS, YMP/PBS). Of all DEGs, 42 genes are differentially expressed in all three treatment groups (that is, increasing with both normal aging and AMP, and decreasing with YMP). b. Bar plot of the top selected biological pathways that are enriched when analyzing either the intersecting DEGs between aging and AMP (149 DEGs) or the intersecting DEGs between aging and YMP (89). Score is derived from GeneAnalytics software. Several pathways affected are shared, suggesting that YMP can reverse some transcriptional consequences of AMP treatment and normal aging. Select genes involved in each pathway are depicted, with DEGs intersecting in all three treatments labeled in red. c. Bar plot of the log fold change of top DEGs that intersect in all three treatments (Aged/Young, AMP/PBS, YMP/PBS). These genes are most likely to be those that are modulated by aged plasma factors in a normal aged milieu, and that this effect can be reversed with exposure to young plasma factors. d. Sankey plot depicting relationships between DEGs which code for BEC surface receptor or membrane proteins, and their corresponding ligands. Directionality of BEC surface protein coding genes in each condition (normal aging, AMP, YMP) are denoted with arrows. Corresponding ligands found significantly up (red) or down (blue) regulated with age in mouse plasma as measured by SOMALogic are highlighted.

To further explore whether the transcriptional changes in response to YMP and AMP may be the result of BEC sensing of peripheral plasma factors, we identified those genes among the 42 intersecting genes (**Figure 6A**) which encoded for receptors or membrane proteins (**Figure 6D**). We then matched the resulting 15 BEC external membrane proteins with putative ligands based on a published resource for receptor-ligand pairs (Ramilowski et al., 2015) and highlighted those ligands which we detected in mouse plasma. Interestingly, we identified Bmpr2 (ligands: BMP7, GDF9), Flt4 (ligand: FN1), Ifnar1 (ligand: IFNA10), Igf1r (ligand: CDH1, GPC3, INS), and Lsr (Ligand: APOB) were not only genes upregulated both with aging and AMP, but some of their ligands were increasing with age in mouse plasma, and have also been reported to increase in human plasma with age (Sun et al., 2018; Tanaka et al., 2018).

## DISCUSSION

Aging is characterized by the gradual decline in physiological integrity and organ function. In the brain, aging is a key risk factor for cognitive decline, neurodegeneration and diseases such as Alzheimer’s disease. With growing evidence that systemic factors, and those in the circulation in particular, can modulate brain aging and function (Wyss-Coray2016), the brain vasculature becomes an obvious putative receiver and transmitter of such circulatory cues to the brain. With this in mind, we characterize here the transcriptome of aging BECs and demonstrate that capillaries are especially sensitive to changes in aging factors in blood.

Single cell RNA sequencing in young mice revealed unique transcriptional signatures for BECs composing arterial, capillary, and venous vascular segments, confirming recent findings by Vanlandewijck and colleagues (Vanlandewijck et al., 2018) and our own lab (Yousef et al., 2019). Importantly, we report here this zonation is not perturbed with age even though significant gene expression changes can be found between ages. Moreover, BEC zonation does not change following systemic exposure of mice to a strong inflammatory trigger, LPS, in spite of several hundred genes changing in unison across the BEC subtypes. Lastly, BEC zonation does not change in mice injected systemically with heterochronic plasma. Together, these observations suggest segmental identity at the transcriptional level is rather stable in response to circulatory environmental cues and possibly defined by more proximal, cellular interactions and signals in the BBB. Additional studies will be necessary to identify these determinants of BEC identity.

While LPS injection induced a concerted upregulation of inflammatory and downregulation of metabolic pathways across all BEC subtypes, aging induced largely zonation specific changes, except a prominent increase in genes involved in translation and RNA biogenesis. Overall, capillary transcriptomes are most responsive to aging as well as intravenous heterochronic plasma administration. Capillary BECs are by far the most abundant segmental subtype in the brain vasculature and while the hemodynamic conditions are different in capillaries than in arterioles and venules, it seems unlikely that this is the cause of their differential response to circulatory cues. It will thus be interesting to determine if capillary BECs are transcriptionally wired to respond to systemic changes and whether this is the result of their interaction with pericytes, other mural cells, glia, and neurons. Equally interesting will be to study the implications of the observed age-related capillary changes on BBB function and neuro-vascular coupling and to endothelial-CNS parenchyma communication in general.

It seems maybe surprising that administration of relatively small amounts of heterochronic plasma (<10% of blood volume per injection) over just 4 days induces robust transcriptional changes in BECs in hundreds of genes. This was particularly evident in capillary BECs of young mice treated with aged plasma (828 DEGs) but also significant in aged mice treated with young plasma (206 DEGs), supporting the notion that capillary BECs are exquisitely and acutely sensitive to changes in the circulation. More importantly, heterochronic plasma injections are sufficient for inducing signatures of aging according to the age of the injected plasma. Aged plasma infusions into young mice strongly induces signatures identified with normal BEC aging including ribosomal RNA processing, hypoxia response, innate immunity, cellular senescence, and TGF-β signaling. The latter may be particularly interesting as increased TGF-β signaling in the vasculature has been linked to age-related basement membrane thickening and cerebral amyloid angiopathy (Wyss-Coray et al., 1997) and to inhibit neural progenitor cell proliferation in the hippocampus (Buckwalter et al., 2006; Yousef et al., 2015a;). Conversely, young plasma administered systemically to aged mice results in a strong downregulation of normal BEC aging signatures including oxidative stress response, innate immunity (via MHC-1), interferon signaling and antigen presentation. *B2m*, which is upregulated by aging and reversed by YMP, is a critical component of major histocompatibility class 1 (MHC-1) molecules (e.g. *H2-K1* and *H2-D1)*, which, enabled by the *Tap1* transporter, allow the recognition of pathogenic antigens by cognate T-cells. These functions can be further augmented by interferons such as *Ifi27* and *Ifnar1*, which is also increased with aging. Not only has soluble *B2m* previously been found at higher levels in human patients with HIV-associated dementia and Alzheimer’s disease (Carrette et al., 2003), it also exerts negative effects on hippocampal neurogenesis and cognition following systemic injection (Smith et al., 2015). Strikingly, young plasma exposure upregulates β-catenin in aged BECs. Wnt/ β-catenin signaling is necessary for maintaining specialized BBB properties, such as tight junction expression and low expression of leukocyte adhesion molecules, but is compromised upon injury, inflammation, and likely during aging (Lengfeld et al., 2017; Liebner et al., 2008; Tran et al., 2015; Zhou and Nathans, 2014). Recently, β-catenin expression alone has been found sufficient to convert leaky vessels in circumventricular organs to a barrier-type state, with stabilized junctions and decreased tracer permeability (Benz et al.; Wang et al., 2019). This suggests factors in young plasma may hold restorative properties for an aged BBB, and could be partly modulated by Wnt/ β-catenin signaling programs.

The 42 genes which mimic an aging transcriptome in young AMP-infused mice and whose expression is reversed (“rejuvenated”) in aged YMP-infused mice are of particular interest. Several of these genes (*Sod1, Apoe, Selenow, Ndufa6, Nostrin*) have established roles in oxidative stress response. Increases in reactive oxygen species (ROS) have been consistently observed in aging and accumulation of oxidative damage to macromolecules is a hallmark of aging, contributing to cellular senescence, loss of proliferation, and secretion of chemokines, interleukins and MMPs (Balaban et al., 2005; Liguori et al., 2018). Interestingly, transcript levels of superoxide dismutase (*Sod1*), an antioxidant shown to increase lifespan, decrease the rate of telomere shortening (Serra et al., 2003) and protect AD model mice (Murakami et al., 2011), are increased in BECs with aging and AMP infusion. Similarly, selenoprotein W *(Selenow*), an antioxidant that protects cells from peroxide-mediated damage (Jeong et al., 2004) and eNOS traffic inducer (*Nostrin)*, an endothelial-specific attenuator of vascular oxidative stress (Förstermann, 2010) are upregulated with aging and AMP infusions, possibly reflecting a protective response induced by factors present in aged plasma. We also find a similar expression pattern in *Rps27* and *Rpl38*, genes involved in ribosomal biogenesis, and *Apoe*, a gene consistently associated with longevity and AD (Kim et al., 2009) and exerting antioxidant properties as well (Jofre-Monseny et al., 2008). Importantly, however, the number of non-intersecting genes between the AMP/normal aging and YMP/normal aging datasets suggests that some aspects of AMP and YMP treatments are not directly antagonistic. YMP treatments can possibly result in the rejuvenation of aging signatures that are not consequences of factors in aged blood but due to other mechanisms of aging. For instance, YMP infusion reduces the expression of genes involved in antigen presentation (*B2m, H2-K1, H2-D1, H2-T23*) and most of these genes are only increased with normal aging and not with AMP infusions. Conversely, expression of regulators of cell death (*Txn, Lmna, Pim1, Ndrg1*) increase with aging and AMP infusion but they are not significantly affected by YMP infusion.

It is likely that many of the changes in BEC gene expression observed with AMP or YMP involve direct receptor-ligand interactions at the luminal surface of these cells. We identified 15 BEC genes - which are not only increased with age but also significantly perturbed by AMP or YMP exposure - that encode for luminal membrane proteins and matched them with their known ligands based on a draft receptor-ligand network in humans (Ramilowski et al., 2015) and our own database of the circulating mouse plasma proteome. Standing out in this list is the IGF1 receptor (Igf1r), which not only increases in expression with aging and aged plasma infusion but its corresponding ligands insulin (INS), glypican 3 (GPC3) and cadherin 1(CDH1) are also increased in aged plasma. Single mutations or deletions of IGF1R homologues increase lifespan in C.elegans (Kimura et al., 1997) and Drosophila (Tatar et al., 2001) and may affect longevity in humans as well (Milman et al., 2016). Other possible mediators of BEC aging are interferons binding to Ifnar1, and BMP or GDFs binding to Bmpr2, all increasing in levels in aged mice plasma and BECs, respectively and linked to aging (Katsimpardi et al., 2014; Loffredo et al., 2013; Yousef et al., 2015a).

## ACKNOWLEDGEMENTS

We thank S. Kolluru and D. Henderson for assistance in library preparation; N. Neff and J. Okamoto for assistance with sequencing; W. Wang, D. Croote, F. Zanini, R. Jones for helpful discussions and computational assistance. This work was funded by the Department of Veterans Affairs (T.W.-C.), the National Institute on Aging (F32-AG051330 to H.Y., R01-AG059694 and DP1-AG053015 to T.W.-C.), the D. H. Chen Foundation (T.W.-C.), The Glenn Foundation for Aging Research (T.W.-C), a SPARK grant to H.Y. through the Stanford Clinical and Translational Science Award (CTSA) to Spectrum (UL1 TR001085) and the Big Idea Brain Rejuvenation Project from the Wu Tsai Neurosciences Institute (T.W.-C.). S.R.Q. is a Chan Zuckerberg Investigator.

## AUTHOR CONTRIBUTIONS

M.B.C., H.Y., A.Y., T.W.-C. designed the research. H.Y. and D.L. performed mouse experiments and provided samples for young/aged healthy mice and AMP treated young mice. A.Y. performed mouse experiments and provided samples for healthy mice and YMP-treated mice. M.B.C. performed single cell library preparation and sequencing pipeline and performed all data analysis, with input from A.Y., HY., and T.W.-C. B.L. and N.S. provided data on the mouse aging plasma proteome. M.B.C. and B.L generated figures. M.B.C., A.Y., T.W.-C. wrote the manuscript with revisions by H.Y and B.L. T.W.-C. and S.R.Q oversaw the project.

## METHODS

### Animals

Aged C57BL6J males were obtained from the National Institute on Aging (NIA), and young C57BL6J males (2-4 months of age) were purchased from Charles River. Mice were housed under a 12-hour light-dark cycle in pathogenic-free conditions, in accordance with the Guide for Care and Use of Laboratory Animals of the National Institutes of Health.

### Plasma collection, dialysis and processing

<u>Mouse:</u> Approximately 500 µl of blood was drawn from the heart in 250 mM EDTA (Sigma Aldrich, CAS Number: <u>60-00-4</u>) and immediately transferred to ice. Blood was centrifuged at 1000g for 10 min at 4°C with a break set to 5 or less. Plasma was collected and immediately snap frozen on dry ice and stored at −80°C until further processing. Plasma was dialyzed in 4L of 1X PBS (51226, Lonza) stirred at room temperature. Plasma was transferred to a fresh 4L of 1 X PBS after 45 min and then again 20 min later. After the second transfer, plasma was dialyzed overnight at 4°C in 4 L of stirred 1X PBS. Plasma from 7-9 mice was pooled for injections.

### Plasma injections in young and aged mice

Young (3-month-old) C57BL6J male mice were treated with 7 injections of aged (18-month-old) or young (3-month-old) dialyzed and pooled mouse plasma (150 uL, r.o.), coming from 8-10 mice per pooled plasma sample as recently described (Yousef et al., 2019). Mice were treated acutely over 4 days, with 2 injections per day spaced 10-12 hours apart. Mice received a 7^th^ plasma injection on day 4 followed by perfusion 3 hours after the last injection. Aged (19-month-old) C57BL6J mice were treated with young (3-month) plasma in the same manner.

### LPS Injections

Mice were injected with Lipopolysaccharide (LPS) derived from Salmonella enterica Serotype Typhimurium (Sigma, L6511), i.p. 1 mg LPS/kg body weight at three successive time points: 0h, 6h, and 24h^85^. Control mice were injected with bodyweight corresponding volumes of PBS. Experimental mice received i.p. and s.c. injections of sterile 0.9% saline with 5% glucose to ensure hydration and stable glucose levels during the procedure. Two hours after the last LPS injection (26h) mouse brains were harvested for BEC isolation and flow analysis.

### Primary BEC isolation and enrichment of CD31+VCAM1+ cells

BEC isolation was based on a previously described procedure(Yousef et al., 2018b). Briefly, mice were anesthetized with avertin and perfused following blood collection. After thoroughly dissecting the meninges, hippocampi were collected, minced and digested using the neural dissociation kit according to kit instructions (Miltenyi, 130-092-628). Brain homogenates were filtered through 35 µm in HBSS and centrifuged pellets were resuspended in 0.9 M sucrose in HBSS followed by centrifugation for 15 min at 850xg at 4°C in order to separate the myelin. This step was repeated for better myelin removal.

Cell pellets were eluted in FACs buffer (0.5% BSA in PBS with 2mM EDTA) and blocked for ten min with Fc preblock (CD16/CD32, BD 553141), followed by 20 minute staining with anti-CD31-APC (1:100, BD 551262), anti-CD45-FITC or anti-CD45-APC/Cy7 (1:100, BD Pharmingen Clone 30-F11 553080; Biolegend, 103116), and anti-Cd11b-BV421 (1:100, Biolegend Clone M1/70 101236). Dead cells were excluded by staining with propidium iodide solution (1:3000, Sigma, P4864). Flow cytometry data and cell sorting were acquired on an ARIA II (BD Biosciences) with FACSDiva software (BD Biosciences). FlowJo software was used for further analysis and depiction of the gating strategy. Gates are indicated by framed areas. Cells were gated on forward (FSC = size) and sideward scatter (SSC = internal structure). FSC-A and FSC-W blotting was used to discriminate single cells from cell doublets/aggregates. PI+ dead cells were excluded. CD11b+ and CD45+ cells were gated to exclude monocytes/macrophages and microglia. CD31+Cd11b-CD45-cells were defined as the BEC population and were sorted directly into lysis buffer in 96 or 384 well plates (Biorad), containing RNAase inhibitor, oligodT, dNTPs and ERCC spike-ins (Picelli et al, 2016), and stored at −80 for further processing. If mice were injected with fluorescently labeled anti-mouse VCAM1-DyLight™488 as described above, CD45 was stained in the APC/Cy7 channel, and CD31+VCAM1+ cells were also gated in the APC and FITC channels.

### Anti-VCAM1 antibody in vivo retro-orbital injections to label CD31+VCAM1+ BECs

Enrichment and gating of VCAM1+ cells was done as previously described(Yousef et al., 2018b). Mice were injected with LPS as described above. When mice received a third LPS injection (24 h), it was followed by retro-orbital injections of either 100µg fluorescently labeled (DyLight™488, Thermo Scientific, 53025) InVivoMAb anti-mouse CD106 (VCAM-1, clone M/K-2.7, Bioxell, BE0027) or fluorescently labeled Rat IgG1 Isotype antibody (Clone HRPN, Bioxell, BE0088). Two hours after the last LPS injection (26h) mouse brains were harvested for BEC isolation and flow analysis.

Healthy young (3-month-old), aged (19-month-old), or plasma injected (r.o.) young mice were similarly injected (r.o.) with fluorescently labeled anti-VCAM1 mAb and gated for flow cytometry analysis of CD31+VCAM1+ cells from hippocampi. Gates are based on positive LPS-stimulated mice injected (r.o.) with anti-VCAM1 or IgG control.

### FACs enrichment of VCAM1 positive BECs

4 young (3-month-old) or 4 aged (19-month-old) C57BL6/J males were injected (r.o.) with fluorescently labeled anti-VCAM1 mAb 2 hours prior to sacrifice and gated for single cell isolation of CD31+VCAM1+ cells from pooled hippocampi following perfusion. Gates are based on positive LPS-stimulated mice injected with fluorescently labeled (DL488) anti-VCAM1 mAb or IgG-DL488 control antibody.

Four hippocampi (from both hemispheres) were pooled together from 4 young (3-month-old) or 4 aged (19-month-old) C57BL6/J males and sorted into lysis buffer in 96-well plates then snap frozen and stored at −80 degrees Celsius until RNA extraction and library preparation. Two, 96-well plates per group contained BECs that were 50% enriched for VCAM1 high expression based on flow cytometry gating; unbiased CD31+ cells were also collected into two, 96-well plates per group.

### Single cell RNA-sequencing

Cell lysis, first-strand synthesis and cDNA synthesis was performed using the Smart-seq-2 protocol as described previously(Picelli et al., 2014) in both 96-well and 384-well formats, with some modifications. After cDNA amplification (23 cycles), cDNA concentrations were determined via capillary electrophoresis (96-well format) or the PicoGreen quantitation assay (384-well format) and wells were cherry-picked to improve quality and cost of sequencing. Cell selection was done through custom scripts and simultaneously normalizes cDNA concentrations to ∼0.2 ng/uL per sample, using the TPPLabtech Mosquito HTS and Mantis (Formulatrix) robotic platforms. Libraries were prepared and pooled using the Illumina Nextera XT kits or and in-house Tn5, following the manufacturer’s instructions. Libraries were then sequenced on the Nextseq or Novaseq (Illumina) using 2 × 75bp paired-end reads and 2 × 8bp index reads with a 200 cycle kit (Illumina, 20012861). Samples were sequenced at an average of 1.5M reads per cell.

### Bioinformatics and data analysis

Sequences from the Nextseq or Novaseq were demultiplexed using bcl2fastq, and reads were aligned to the mm10 genome augmented with ERCC sequences, using STAR version 2.5.2b. Gene counts were made using HTSEQ version 0.6.1p1. All packages were called an run through a custom Snakemake pipeline. We applied standard algorithms for cell filtration, feature selection, and dimensionality reduction. Briefly, genes appearing in fewer than 5 cells, samples with fewer than 100 genes, and samples with less than 50,000 reads were excluded from the analysis. Out of these cells, those with more than 30% of reads as ERCC, and more than 10% mitochondrial or 10% ribosomal were also excluded from analysis. Counts were log-normalized then scaled.

Next, the *Canonical Correlation Analysis* function from the Seurat package (Butler et al., 2018) was used to align raw data from multiple experiments, data from aged vs young mice, AMP vs YMP treated young mice, and LPS treated vs untreated mice. Only the first 10 Canonical Components (CCs) were used. After alignment, relevant features were selected by filtering expressed genes to a set of ∼3000 with the highest positive and negative pairwise correlations. Genes were then projected into principal component space using the robust principal component analysis (rPCA). Single cell PC scores and genes loadings for the first 20 PCs were used as inputs into Seurat’s (v2) *FindClusters* and *RunTsne* functions to calculate 2-dimensional tSNE coordinates and search for distinct cell populations. Briefly, a shared-nearest-neighbor graph was constructed based on the Euclidean distance metric in PC space, and cells were clustered using the Louvain method. Cells and clusters were then visualized using 3-D t-distributed Stochastic Neighbor embedding on the same distance metric. Differential gene expression analysis was done by applying the Mann-Whitney U-test of the BEC clusters obtained using unsupervised clustering. Raw p-values were adjusted via the false discovery rate (FDR). Permutation tests were then performed on all genes of interest. All graphs and analyses were generated and performed in R. GeneAnalytics and GeneCards-packages offered by Gene Set Enrichment Analysis (GSEA) tool was used for GO/KEGG/REACTOME pathway analysis and classification of enriched genes in each subpopulation.

## REFERENCES

Abbott, N.J., Rönnbäck, L., and Hansson, E. (2006). Astrocyte-endothelial interactions at the blood-brain barrier. Nat. Rev. Neurosci. 7, 41–53.

Andreone, B.J., Chow, B.W., Tata, A., Lacoste, B., Ben-Zvi, A., Bullock, K., Deik, A.A., Ginty, D.D., Clish, C.B., and Gu, C. (2017). Blood-Brain Barrier Permeability Is Regulated by Lipid Transport-Dependent Suppression of Caveolae-Mediated Transcytosis. Neuron 94, 581–594.e5.

Andrews-Hanna, J.R., Snyder, A.Z., Vincent, J.L., Lustig, C., Head, D., Raichle, M.E., and Buckner, R.L. (2007). Disruption of Large-Scale Brain Systems in Advanced Aging. Neuron 56, 924–935.

Armulik, A., Genové, G., Mäe, M., Nisancioglu, M.H., Wallgard, E., Niaudet, C., He, L., Norlin, J., Lindblom, P., Strittmatter, K., et al. (2010). Pericytes regulate the blood-brain barrier. Nature 468, 557–561.

Balaban, R.S., Nemoto, S., and Finkel, T. (2005). Mitochondria, Oxidants, and Aging. Cell 120, 483–495.

Ben-Zvi, A., Lacoste, B., Kur, E., Andreone, B.J., Mayshar, Y., Yan, H., and Gu, C. (2014). Mfsd2a is critical for the formation and function of the blood-brain barrier. Nature 509, 507–511.

Benz, F., Wichitnaowarat, V., Lehmann, M., and Germano, R.F. V Low Wnt / β - catenin signaling determines leaky vessels in the subfornical organ and affects water homeostasis in mice.

Bien-Ly, N., Boswell, C.A., Jeet, S., Beach, T.G., Hoyte, K., Luk, W., Shihadeh, V., Ulufatu, S., Foreman, O., Lu, Y., et al. (2015). Lack of Widespread BBB Disruption in Alzheimer’s Disease Models: Focus on Therapeutic Antibodies. Neuron 88, 289–297.

Bishop, N.A., Lu, T., and Yankner, B.A. (2010). Neural mechanisms of ageing and cognitive decline. Nature 464, 529–535.

Broadwell, R.D. (1989). Transcytosis of macromolecules through the blood-brain barrier: a cell biological perspective and critical appraisal. Acta Neuropathol. 79, 117–128.

Buckwalter, M.S., Yamane, M., Coleman, B.S., Ormerod, B.K., Chin, J.T., Palmer, T., and Wyss-Coray, T. (2006). Chronically increased transforming growth factor-beta1 strongly inhibits hippocampal neurogenesis in aged mice. Am. J. Pathol. 169, 154–164.

Butler, A., Hoffman, P., Smibert, P., Papalexi, E., and Satija, R. (2018). Analysis Integrating single-cell transcriptomic data across different conditions, technologies, and species. Nat. Biotechnol. 36, 411–420.

Carrette, O., Demalte, I., Scherl, A., Yalkinoglu, O., Corthals, G., Burkhard, P., Hochstrasser, D.F., and Sanchez, J.-C. (2003). A panel of cerebrospinal fluid potential biomarkers for the diagnosis of Alzheimer’s disease. Proteomics 3, 1486–1494.

Castellano, J.M., Mosher, K.I., Hinkson, I. V., Tingle, M., Angst, M.S., Zou, B., Berdnik, D., McBride, A.A., Shen, J.C., Xie, X.S., et al. (2017). Human umbilical cord plasma proteins revitalize hippocampal function in aged mice. Nature 544, 488–492.

Chow, B.W., and Gu, C. (2015). The Molecular Constituents of the Blood-Brain Barrier. Trends Neurosci. 38, 598–608.

Daneman, R., and Prat, A. (2014). The Blood Brain Barrier (BBB). Cold Spring Harb. Perspect. Biol. 10, 1–24.

Daneman, R., Zhou, L., Kebede, A.A., and Barres, B.A. (2010). Pericytes are required for bloodg€”brain barrier integrity during embryogenesis. Nature 468, 562–566.

Förstermann, U. (2010). Nitric oxide and oxidative stress in vascular disease. Pflügers Arch. - Eur. J. Physiol. 459, 923–939.

Iadecola, C. (2013). The Pathobiology of Vascular Dementia. Neuron 80, 844–866.

Jeong, D., Kim, E.H., Kim, T.S., Chung, Y.W., Kim, H., and Kim, I.Y. (2004). Different Distributions of Selenoprotein W and Thioredoxin during Postnatal Brain Development and Embryogenesis. Mol. Cells 17, 156–159.

Jofre-Monseny, L., Minihane, A.-M., and Rimbach, G. (2008). Impact of apoE genotype on oxidative stress, inflammation and disease risk. Mol. Nutr. Food Res. 52, 131–145.

Katsimpardi, L., Litterman, N.K., Schein, P.A., Miller, C.M., Loffredo, F.S., Wagers, A.J., Lee, R.T., Chen, J.W., Wojtkiewicz, G.R., and Rubin, L.L. (2014). Vascular and Neurogenic Rejuvenation of the Aging Mouse Brain by Young Systemic Factors. Science (80-.). 344, 630–634.

Khrimian, L., Obri, A., Ramos-Brossier, M., Rousseaud, A., Moriceau, S., Nicot, A.-S., Karsenty, G., Gao, X.-B., Oury, F., Kosmidis, S., et al. (2017). Gpr158 mediates osteocalcin’s regulation of cognition. J. Exp. Med. 214, 2859–2873.

Kim, J., Basak, J.M., and Holtzman, D.M. (2009). The role of apolipoprotein E in Alzheimer’s disease. Neuron 63, 287–303.

Kimura, K.D., Tissenbaum, H.A., Liu, Y., and Ruvkun, G. (1997). Daf-2, an Insulin Receptor-Like Gene That Regulates Longevity and Diapause in Caenorhabditis elegans. Science (80-.). 277, 942 LP–946.

Lengfeld, J.E., Lutz, S.E., Smith, J.R., Diaconu, C., Scott, C., Kofman, S.B., Choi, C., Walsh, C.M., Raine, C.S., Agalliu, I., et al. (2017). Endothelial Wnt / β -catenin signaling reduces immune cell infiltration in multiple sclerosis.

Liebner, S., Corada, M., Bangsow, T., Babbage, J., Taddei, A., Czupalla, C.J., Reis, M., Felici, A., Wolburg, H., Fruttiger, M., et al. (2008). Wnt/ beta-catenin signaling controls development of the blood –brain barrier. 183, 409–417.

Liguori, I., Russo, G., Curcio, F., Bulli, G., Aran, L., Della-Morte, D., Gargiulo, G., Testa, G., Cacciatore, F., Bonaduce, D., et al. (2018). Oxidative stress, aging, and diseases. Clin. Interv. Aging 13, 757–772.

Loffredo, F.S., Steinhauser, M.L., Jay, S.M., Gannon, J., Pancoast, J.R., Yalamanchi, P., Sinha, M., Dall’Osso, C., Khong, D., Shadrach, J.L., et al. (2013). Growth Differentiation Factor 11 Is a Circulating Factor that Reverses Age-Related Cardiac Hypertrophy. Cell 153, 828–839.

López-Otín, C., Blasco, M.A., Partridge, L., Serrano, M., and Kroemer, G. (2013). The hallmarks of aging. Cell 153.

Marques, F., Sousa, J.C., Sousa, N., and Palha, J.A. (2013). Blood –brain-barriers in aging and in Alzheimer‘s disease. Mol. Neurodegener. 8, 1–9.

Mattson, M.P., and Magnus, T. (2006). Ageing and neuronal vulnerability. Nat. Rev. Neurosci. 7, 278–294.

Milman, S., Huffman, D.M., and Barzilai, N. (2016). The Somatotropic Axis in Human Aging: Framework for the Current State of Knowledge and Future Research. Cell Metab. 23, 980–989.

Montagne, A., Barnes, S.R., Sweeney, M.D., Halliday, M.R., Sagare, A.P., Zhao, Z., Toga, A.W., Jacobs, R.E., Liu, C.Y., Amezcua, L., et al. (2015). Blood-Brain barrier breakdown in the aging human hippocampus. Neuron 85, 296–302.

Mooradian, A.D. (1988). Effect of aging on the blood-brain barrier. Neurobiol. Aging 9, 31–39.

Murakami, K., Murata, N., Noda, Y., Tahara, S., Kaneko, T., Kinoshita, N., Hatsuta, H., Murayama, S., Barnham, K.J., Irie, K., et al. (2011). SOD1 (copper/zinc superoxide dismutase) deficiency drives amyloid β protein oligomerization and memory loss in mouse model of Alzheimer disease. J. Biol. Chem. 286, 44557–44568.

Picelli, S., Faridani, O.R., Björklund, Å.K., Winberg, G., Sagasser, S., and Sandberg, R. (2014). Full-length RNA-seq from single cells using Smart-seq2. Nat. Protoc. 9, 171–181.

Ramilowski, J.A., Goldberg, T., Harshbarger, J., Kloppmann, E., Lizio, M., Satagopam, V.P., Itoh, M., Kawaji, H., Carninci, P., Rost, B., et al. (2015). A draft network of ligand–receptormediated multicellular signalling in human. Nat. Commun. 6, 7866.

Reese, T.S., and Karnovsky, M.J. (2004). Fine Structural Localization of a Blood-Brain Barrier To Exogenous Peroxidase. J. Cell Biol. 34, 207–217.

Serra, V., von Zglinicki, T., Lorenz, M., and Saretzki, G. (2003). Extracellular Superoxide Dismutase Is a Major Antioxidant in Human Fibroblasts and Slows Telomere Shortening. J. Biol. Chem. 278, 6824–6830.

Silva-Vargas, V., Maldonado-Soto, A.R., Mizrak, D., Codega, P., and Doetsch, F. (2016). Age-Dependent Niche Signals from the Choroid Plexus Regulate Adult Neural Stem Cells. Cell Stem Cell 19, 643–652.

Smith, L.K., He, Y., Park, J.S., Bieri, G., Snethlage, C.E., Lin, K., Gontier, G., Wabl, R., Plambeck, K.E., Udeochu, J., et al. (2015). B2-Microglobulin Is a Systemic Pro-Aging Factor That Impairs Cognitive Function and Neurogenesis. Nat. Med. 21, 932–937.

Sun, B.B., Maranville, J.C., Peters, J.E., Stacey, D., Surendran, P., Danesh, J., Roberts, D.J., Fox, C.S., Oliver-Williams, C., Staley, J.R., et al. (2018). Genomic atlas of the human plasma proteome. Nature 558, 73–79.

Sweeney, M.D., Kisler, K., Montagne, A., Toga, A.W., and Zlokovic, B. V. (2018). The role of brain vasculature in neurodegenerative disorders. Nat. Neurosci. 21, 1318–1331.

Tanaka, T., Biancotto, A., Moaddel, R., Moore, A.Z., Gonzalez-Freire, M., Aon, M.A., Candia, J., Zhang, P., Cheung, F., Fantoni, G., et al. (2018). Plasma proteomic signature of age in healthy humans. Aging Cell 17, 1–13.

Tatar, M., Kopelman, A., Epstein, D., Tu, M.-P., Yin, C.-M., and Garofalo, R.S. (2001). A Mutant Drosophila Insulin Receptor Homolog That Extends Life-Span and Impairs Neuroendocrine Function. Science (80-.). 292, 107 LP–110.

Tran, K.A., Zhang, X., Predescu, D., and Huang, X. (2015). Endothelial β -Catenin Signaling Is Required for Maintaining Adult Blood –Brain Barrier Integrity and Central Nervous. 177–186.

Vanlandewijck, M., He, L., Mäe, M.A., Andrae, J., Ando, K., Del Gaudio, F., Nahar, K., Lebouvier, T., Laviña, B., Gouveia, L., et al. (2018). A molecular atlas of cell types and zonation in the brain vasculature. Nature 554, 475–480.

Villeda, S.A., Luo, J., Mosher, K.I., Zou, B., Britschgi, M., Bieri, G., Stan, T.M., Fainberg, N., Ding, Z., Eggel, A., et al. (2011). The ageing systemic milieu negatively regulates neurogenesis and cognitive function. Nature 477, 90–96.

Villeda, S.A., Plambeck, K.E., Middeldorp, J., Castellano, J.M., Mosher, K.I., Luo, J., Smith, L.K., Bieri, G., Lin, K., Berdnik, D., et al. (2014). Young blood reverses age-related impairments in cognitive function and synaptic plasticity in mice. Nat. Med. 20, 659–663.

Wang, Y., Sabbagh, M.F., Gu, X., and Rattner, A. (2019). Beta-catenin signaling regulates barrierspecific gene expression in circumventricular organ and ocular vasculatures. 1–36.

Wyss-Coray, T. (2016). Ageing, neurodegeneration and brain rejuvenation. Nature 539, 180–186.

Wyss-Coray, T., Masliah, E., Mallory, M., McConlogue, L., Johnson-Wood, K., Lin, C., and Mucke, L. (1997). Amyloidogenic role of cytokine TGF-β1 in transgenic mice and in Alzheimer’s disease. Nature 389, 603.

Yousef, H., Morgenthaler, A., Schlesinger, C., Bugaj, L., Conboy, I.M., and Schaffer, D. V (2015a). Age-Associated Increase in BMP Signaling Inhibits Hippocampal Neurogenesis. Stem Cells 33, 1577–1588.

Yousef, H., Conboy, M.J., Morgenthaler, A., Schlesinger, C., Bugaj, L., Paliwal, P., Greer, C., Conboy, I.M., Schaffer, D., Yousef, H., et al. (2015b). Systemic attenuation of the TGF-β pathway by a single drug simultaneously rejuvenates hippocampal neurogenesis and myogenesis in the same old mammal. Oncotarget 6, 11959–11978.

Yousef, H., Czupalla, C.J., Lee, D., Burke, A., Chen, M., Zandstra, J., Berber, E., Lehallier, B., Mathur, V., Nair, R. V, et al. (2018a). Aged blood inhibits hippocampal neurogenesis and activates microglia through VCAM1 at the blood-brain barrier. BioRxiv.

Yousef, H., Czupalla, C., Wyss-Coray, T., and Butcher, E. (2018b). Papain-based Single Cell Isolation of Primary Murine Brain Endothelial Cells Using Flow Cytometry. Bio-Protocol 8, 1–12.

Zhao, Z., Nelson, A.R., Betsholtz, C., and Zlokovic, B. V. (2015). Establishment and Dysfunction of the Blood-Brain Barrier. Cell 163, 1064–1078.

Zhou, Y., and Nathans, J. (2014). Short Article Gpr124 Controls CNS Angiogenesis and Blood-Brain Barrier Integrity by Promoting. Dev. Cell 31, 248–256.

Zlokovic, B. V. (2008). The Blood-Brain Barrier in Health and Chronic Neurodegenerative Disorders. Neuron 57, 178–201.

