## Supplementary Information for "Brain endothelial cells are exquisite sensors of age-related circulatory cues"

**SUPPLEMENTARY FIGURE LEGENDS**

**Figure SI 1**

a. Barplots of the number of reads for each cell (pre-QC) that is not aligned, not unique and properly aligned, for all 4 replicates of young and aged cells (total 2034 cells).

b. Scatter-plot of the number of reads vs the number of genes expressed per cell (pre-QC). Cells are then filtered to those expressing at east 500 genes or at least 50, 000 aligned (mapped) reads.

c. Number of cells per biological replicate of aged and young cells that remain after QC is applied.

d. Violin plots of the number of mapped reads per cell within each of the biological replicates for aged and young cells combined.

e. Violin plots of the number of genes expressed per cell within each of the biological replicates for aged and young cells combined.

f. Scatterplot of the percentage of genes expressed per cell (by age) that is mitochondrial or ribosomal.

**Figure SI 2**

a. Ratio of all sorted cells (based only on CD31+/CD45 -) that identifies as either arterial (A) or venous (V) in segmental identity (based on the expression of at least ⅓ canonical marker genes), for both aged and young mice. Note the break in the axes.

b. Scatter-plot of the spearman's correlation with Vcam1 in either the arterial or venous populations. Note the high correlation of Vcam1 with canonical arterial markers Vegfc, Bmx, Gja4 and venous markers Nr2f2, Slc38a5.

c. Volcano plots of the top DEGs between *Vcam1*+ and *Vcam1*- cells in either arterial or venous populations. Note the transcriptional activation of *Vcam1*+ cells in both populations. Dotted heatmaps show the average expression level of key arterial and venous defining markers and the relative differences in percent expression in *Vcam1*+ and Vcam1- populations.

Ideal arterial and venous markers should not be differentially expressed between *Vcam1*+/- populations, in order to best encompass the range of arterial and venous cells.

d. The total number of cells collected from each replicate that identifies as A,C or V, based on unbiased transcriptome clustering, after the addition of *Vcam1* enriched samples (~20% of all cells).

**Figure SI 3**

a. Ratio of A-C-V cells collected in LPS-treated and untreated mice through unbiased CD31+/CD45- sorting remains largely unchanged.

b. Scatterplot showing the genes which are commonly and oppositely differentially expressed (FDR<0.1 in both) between aging capillaries, and LPS treated capillaries. Notes the relatively low number of commonly upregulated DEGs (blue).

**Figure SI 4**

Pairwise plot of all 2034 young and aged BECs over the first 10 aligned correlation components (ACC). Cells are colored by their defined segmental identity obtained from running SNN clustering on the first 10 ACCs. Note that the first 3 ACCs largely define A-C-V identities.

**Figure SI 5**

Pairwise plot of all 2034 young and aged BECs over the first 10 aligned correlation components (ACC). Cells are colored by their age group. Note that no ACCs appear to explain heterogeneities between the two age groups.

**Figure SI 6**

a. tSNE plot of all aged and young BECs together, colored by number of genes (left) and number of reads mapped (right).

b. Violin plots of the number of genes expressed per cell, in aged and young groups.

c. Violin plots of the number of genes expressed per cell, in A-C-V groups.

d. Violin plot of the permuted distribution (when cell age labels are shuffled) of average log fold changes for several genes of interest. All genes show true observed values well above the 95th percentile of the permuted distribution.

**Figure SI 7**

a. Ratio of A-C-V cells collected in AMP and PBS treated mice through unbiased CD31+/CD45- sorting remains largely unchanged.

b. Scatterplot showing the genes which are commonly and oppositely differentially expressed (FDR<0.1 in both) between aging and AMP treated venous cells. Note the relatively low number of common DEGs (red).

c. Scatterplot showing the genes which are commonly and oppositely differentially expressed (FDR<0.1 in both) between aging and AMP treated arterial cells. Note the relatively low number of common DEGs (red).

d. GO analysis of the pathways enriched in the genes upregulated and downregulated with AMP treatment (compared to PBS)

e. Scatter-plot of the -log10(FDR)*log2FC values for each gene. Genes differentially up and down regulated are marked in red. Top scoring genes are further labeled.

f. Scatter-plot of the mean log2CPM values of all genes in either AMP treated or aged BECs. Common DEGs between the two conditions (b) are depicted in red. Top expressed genes are labeled in blue. Note that all common DEGs are more highly expressed in AMP treatment than disease-free aging.

g. Violin plot of the permuted distribution (when cell treatment labels are shuffled) of average log fold changes for several genes of interest in capillary cells, and the relative positions of the true observations and the 95th percentile of the permuted distributions.

**Fig SI 8**

a. GO analysis of the pathways enriched in the genes upregulated and downregulated with YMP treatment (compared to PBS)

b. Scatter-plot of the -log10(FDR)*log2FC values for each gene. Genes differentially up and down regulated are marked in red. Top scoring genes are further labeled.
c. Scatter-plot of the mean log2CPM values of all genes in either YMP treated or aged BECs. Common DEGs between the two conditions (b) are depicted in red. Top expressed genes are labeled in blue.
d. Violin plot of the permuted distribution (when cell treatment labels are shuffled) of average log fold changes for several genes of interest in capillary cells, and the relative positions of the true observations and the 95th percentile of the permuted distributions.

**Fig SI 9**

Bubble plots of key genes grouped by function, and their log fold change values in each treatment (Aged/Young, YMP/PBS, AMP/PBS). Gene of interest are highlighted in grey.

**Supplementary Tables**

**SI Table 1**

List of genes that are commonly differentially up- or down-regulated in both AMP (in young) v.s. PBS (in young) as well as YMP (in young) vs PBS (in young).

*Figure SI 1*

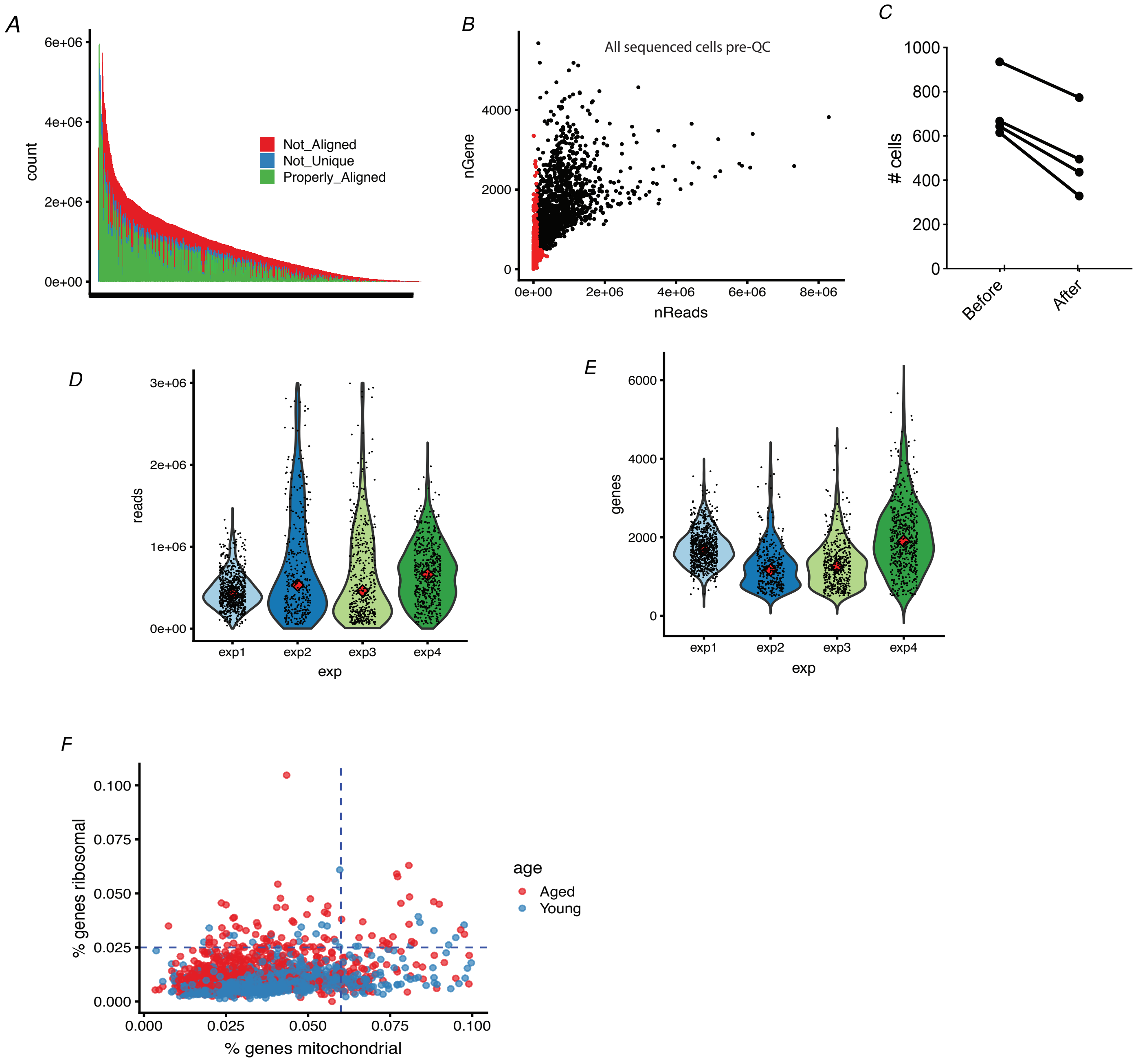
A
6e+06
4e+06
2e+06
0e+00
count
Not_Aligned
Not_Unique
Properly_Aligned
B
All sequenced cells pre-QC
nGene
4000
2000
0
0e+00
2e+06
4e+06
6e+06
8e+06
nReads
C
1000
800
600
400
200
0
### cells
Before
After
D
3e+06
2e+06
1e+06
0e+00
reads
exp1
exp2
exp3
exp4
exp
E
6000
4000
2000
0
genes
exp1
exp2
exp3
exp4
exp
F
0.100
0.075
0.050
0.025
0.000
% genes ribosomal
0.000
0.025
0.050
0.075
0.100
% genes mitochondrial
age
Aged
Young

*Figure SI 2*

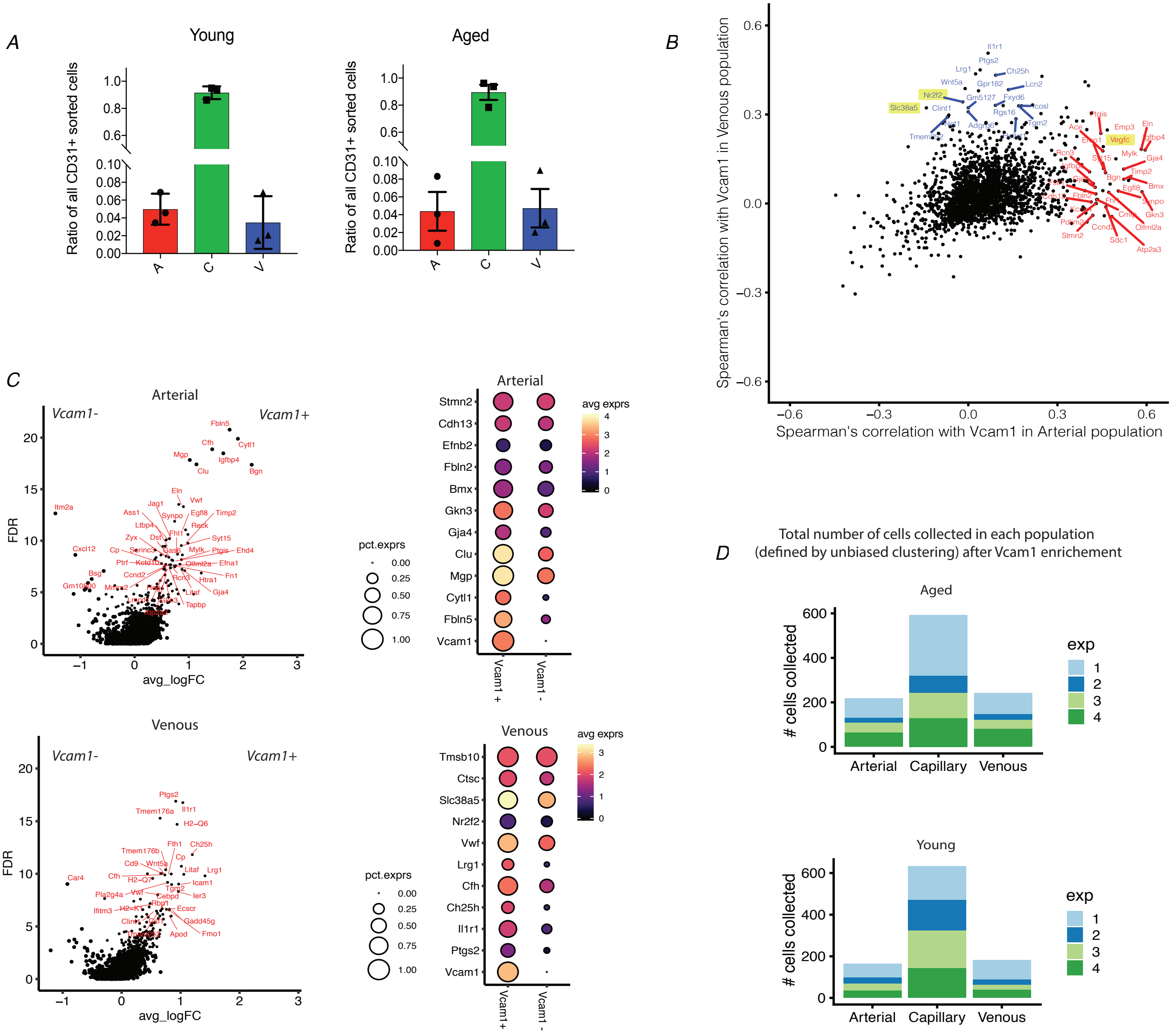
A
Young
Aged
Ratio of all CD31+ sorted cells
A
C
V
B
Spearman's correlation with Vcam1 in Venous population
Spearman's correlation with Vcam1 in Arterial population
C
Arterial
Vcam1-
Vcam1+
FDR
avg_logFC
pct.exprs
avg exprs
Stmn2
Cdh13
Efnb2
Fbln2
Bmx
Gkn3
Gja4
Clu
Mgp
Cytl1
Fbln5
Vcam1
Vcam1 +
Vcam1 -
Venous
Tmsb10
Ctsc
Slc38a5
Nr2f2
Vwf
Lrg1
Cfh
Ch25h
Il1r1
Ptgs2
D
Total number of cells collected in each population
(defined by unbiased clustering) after Vcam1 enrichement
Aged
Young
### cells collected
exp
Arterial
Capillary
Venous

*Figure SI 3*

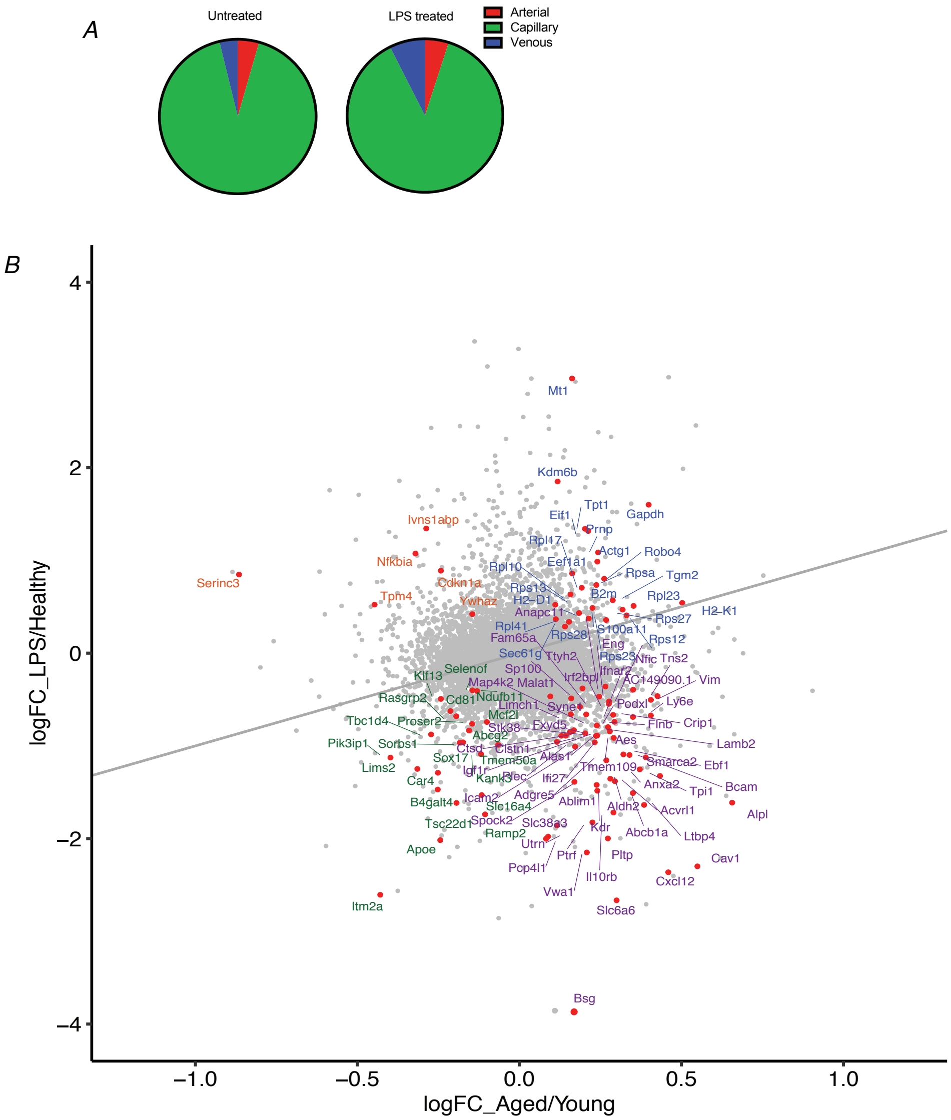
A
Untreated
LPS treated
Arterial
Capillary
Venous
B
logFC_LPS/Healthy
logFC_Aged/Young
4
2
0
−2
−4
−1.0
−0.5
0.0
0.5
1.0
Mt1
Kdm6b
Gapdh
Tpt1
Eif1
Prnp
Ivns1abp
Nfkbia
Serinc3
Cdkn1a
Tpm4
Ywhaz
Bsg
Itm2a
Apoe
Vim
Cav1
Cxcl12
Slc6a6
Alpl
Bcam
Lamb2

*Figure SI 4*

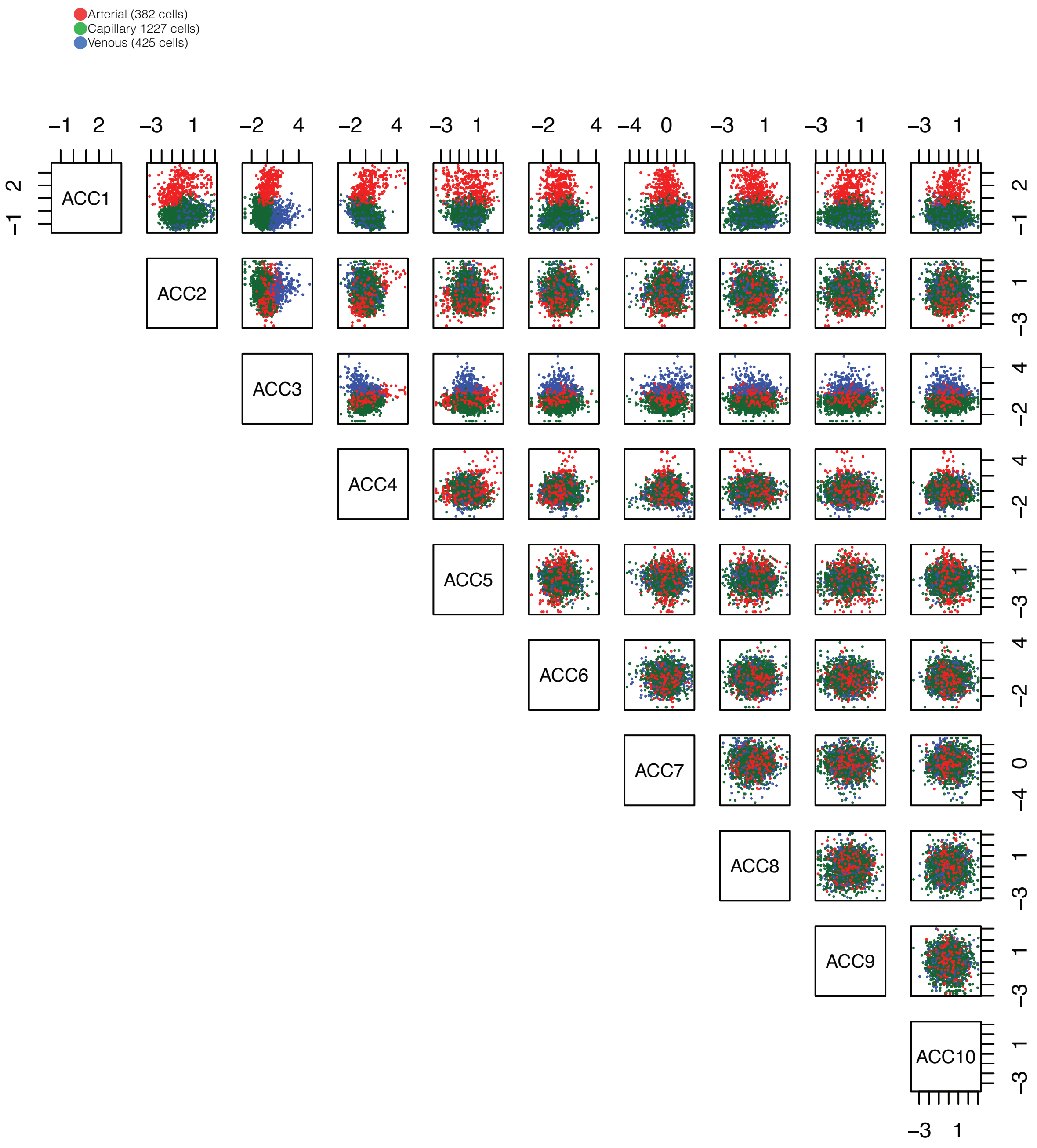

Arterial (382 cells)
Capillary 1227 cells)
Venous (425 cells)
ACC1
ACC2
ACC3
ACC4
ACC5
ACC6
ACC7
ACC8
ACC9
ACC10

*Figure SI 5*

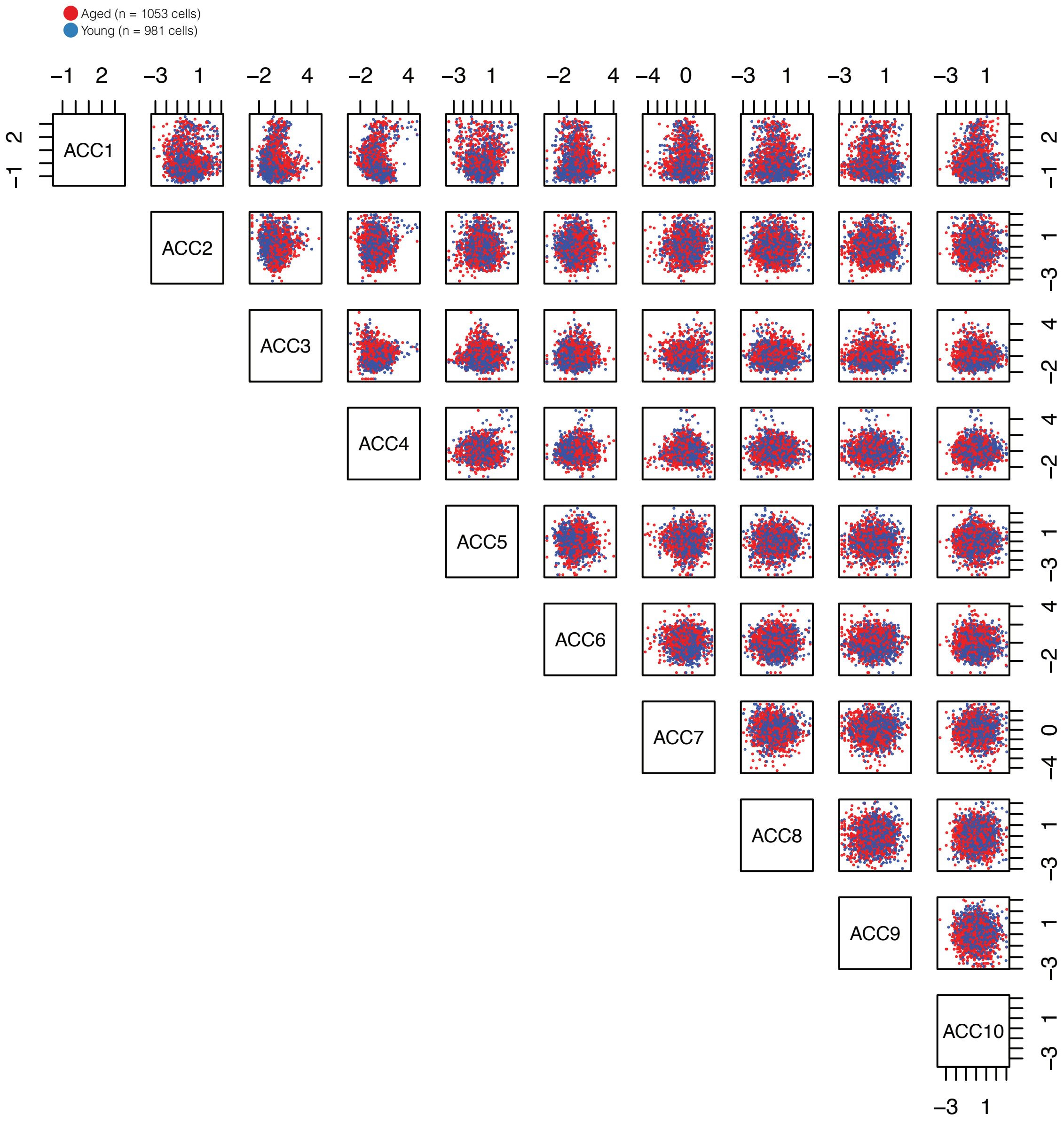

Aged (n = 1053 cells)
Young (n = 981 cells)
ACC1
ACC2
ACC3
ACC4
ACC5
ACC6
ACC7
ACC8
ACC9
ACC10

*Figure SI 6*

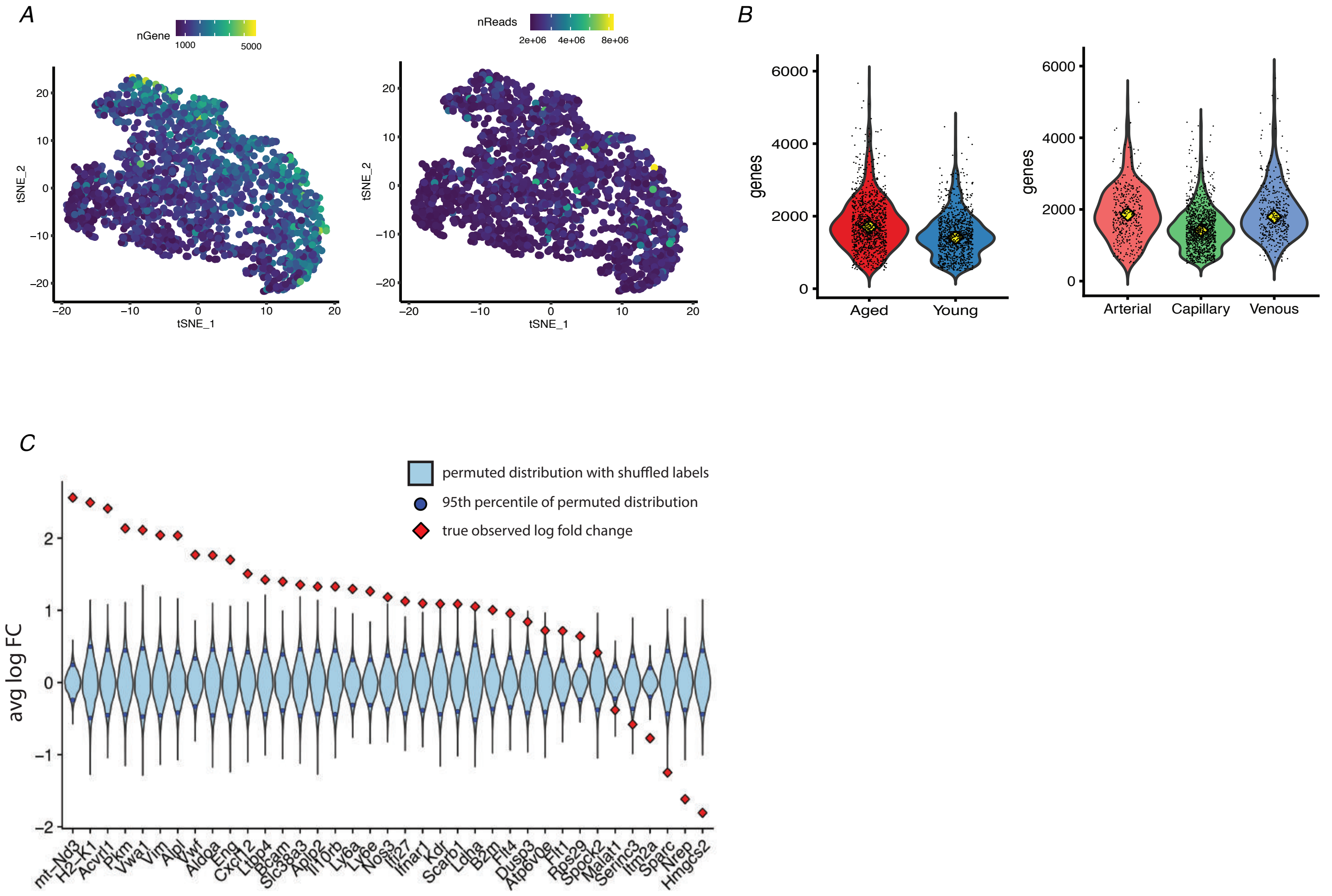
A
nGene
1000
5000
nReads
2e+06
4e+06
8e+06
tSNE_1
tSNE_2
B
genes
Aged
Young
Arterial
Capillary
Venous
C
permuted distribution with shuffled labels
95th percentile of permuted distribution
true observed log fold change
avg log FC
mt–Nd3
H2–K1
Acvrl1
Pkm
Vwa1
Vim
Alpl
Vwf
Aldoa
Eng
Cxcl12
Ltbp4
Bcam
Slc38a3
Aplp2
Il10rb
Ly6a
Ly6e
Nos3
Ifi27
Ifnar1
Kdr
Scarb1
Ldha
B2m
Flt4
Dusp3
Atp6v0e
Flt1
Rps29
Spock2
Malat1
Serinc3
Itm2a
Sparc
Nrep
Hmgcs2

SI 7

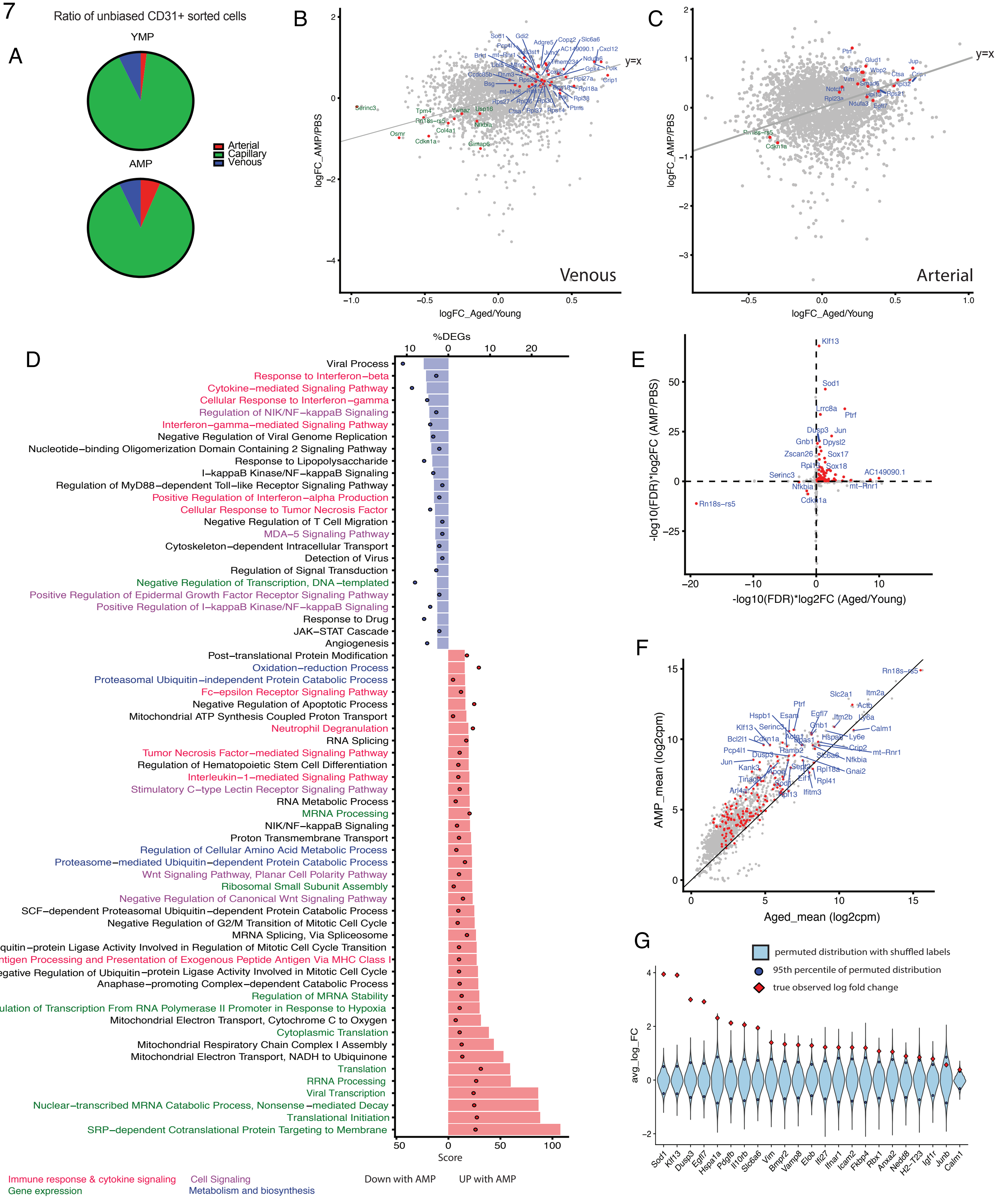
A
Ratio of unbiased CD31+ sorted cells
YMP
AMP
Arterial
Capillary
Venous
B
logFC_AMP/PBS
logFC_Aged/Young
y=x
Venous
C
logFC_AMP/PBS
logFC_Aged/Young
y=x
Arterial
D
%DEGs
Viral Process
Response to Interferon−beta
Cytokine−mediated Signaling Pathway
Cellular Response to Interferon−gamma
Regulation of NIK/NF−kappaB Signaling
Interferon−gamma−mediated Signaling Pathway
Negative Regulation of Viral Genome Replication
Nucleotide−binding Oligomerization Domain Containing 2 Signaling Pathway
Response to Lipopolysaccharide
I−kappaB Kinase/NF−kappaB Signaling
Regulation of MyD88−dependent Toll−like Receptor Signaling Pathway
Positive Regulation of Interferon−alpha Production
Cellular Response to Tumor Necrosis Factor
Negative Regulation of T Cell Migration
MDA−5 Signaling Pathway
Cytoskeleton−dependent Intracellular Transport
Detection of Virus
Regulation of Signal Transduction
Negative Regulation of Transcription, DNA−templated
Positive Regulation of Epidermal Growth Factor Receptor Signaling Pathway
Positive Regulation of I−kappaB Kinase/NF−kappaB Signaling
Response to Drug
JAK−STAT Cascade
Angiogenesis
Post−translational Protein Modification
Oxidation−reduction Process
Proteasomal Ubiquitin−independent Protein Catabolic Process
Fc−epsilon Receptor Signaling Pathway
Negative Regulation of Apoptotic Process
Mitochondrial ATP Synthesis Coupled Proton Transport
Neutrophil Degranulation
RNA Splicing
Tumor Necrosis Factor−mediated Signaling Pathway
Regulation of Hematopoietic Stem Cell Differentiation
Interleukin−1−mediated Signaling Pathway
Stimulatory C−type Lectin Receptor Signaling Pathway
RNA Metabolic Process
MRNA Processing
NIK/NF−kappaB Signaling
Proton Transmembrane Transport
Regulation of Cellular Amino Acid Metabolic Process
Proteasome−mediated Ubiquitin−dependent Protein Catabolic Process
Wnt Signaling Pathway, Planar Cell Polarity Pathway
Ribosomal Small Subunit Assembly
Negative Regulation of Canonical Wnt Signaling Pathway
SCF−dependent Proteasomal Ubiquitin−dependent Protein Catabolic Process
Negative Regulation of G2/M Transition of Mitotic Cell Cycle
MRNA Splicing, Via Spliceosome
Ubiquitin−protein Ligase Activity Involved in Regulation of Mitotic Cell Cycle Transition
Antigen Processing and Presentation of Exogenous Peptide Antigen Via MHC Class I
Negative Regulation of Ubiquitin−protein Ligase Activity Involved in Mitotic Cell Cycle
Anaphase−promoting Complex−dependent Catabolic Process
Regulation of MRNA Stability
Regulation of Transcription From RNA Polymerase II Promoter in Response to Hypoxia
Mitochondrial Electron Transport, Cytochrome C to Oxygen
Cytoplasmic Translation
Mitochondrial Respiratory Chain Complex I Assembly
Mitochondrial Electron Transport, NADH to Ubiquinone
Translation
RRNA Processing
Viral Transcription
Nuclear−transcribed MRNA Catabolic Process, Nonsense−mediated Decay
Translational Initiation
SRP−dependent Cotranslational Protein Targeting to Membrane
Score
Immune response & cytokine signaling
Cell Signaling
Gene expression
Metabolism and biosynthesis
Down with AMP
UP with AMP
E
-log10(FDR)*log2FC (AMP/PBS)
-log10(FDR)*log2FC (Aged/Young)
F
AMP_mean (log2cpm)
Aged_mean (log2cpm)
G
permuted distribution with shuffled labels
95th percentile of permuted distribution
true observed log fold change
avg_log_FC
Sod1
Klf13
Dusp3
Egfl7
Hspa1a
Pdgfb
Il10rb
Slc6a6
Vim
Bmpr2
Vamp8
Elob
Ifi27
Ifnar1
Icam2
Fkbp4
Rbx1
Anxa2
Nedd8
H2-T23
Igf1r
Junb
Calm1

Figure SI 8: YMP

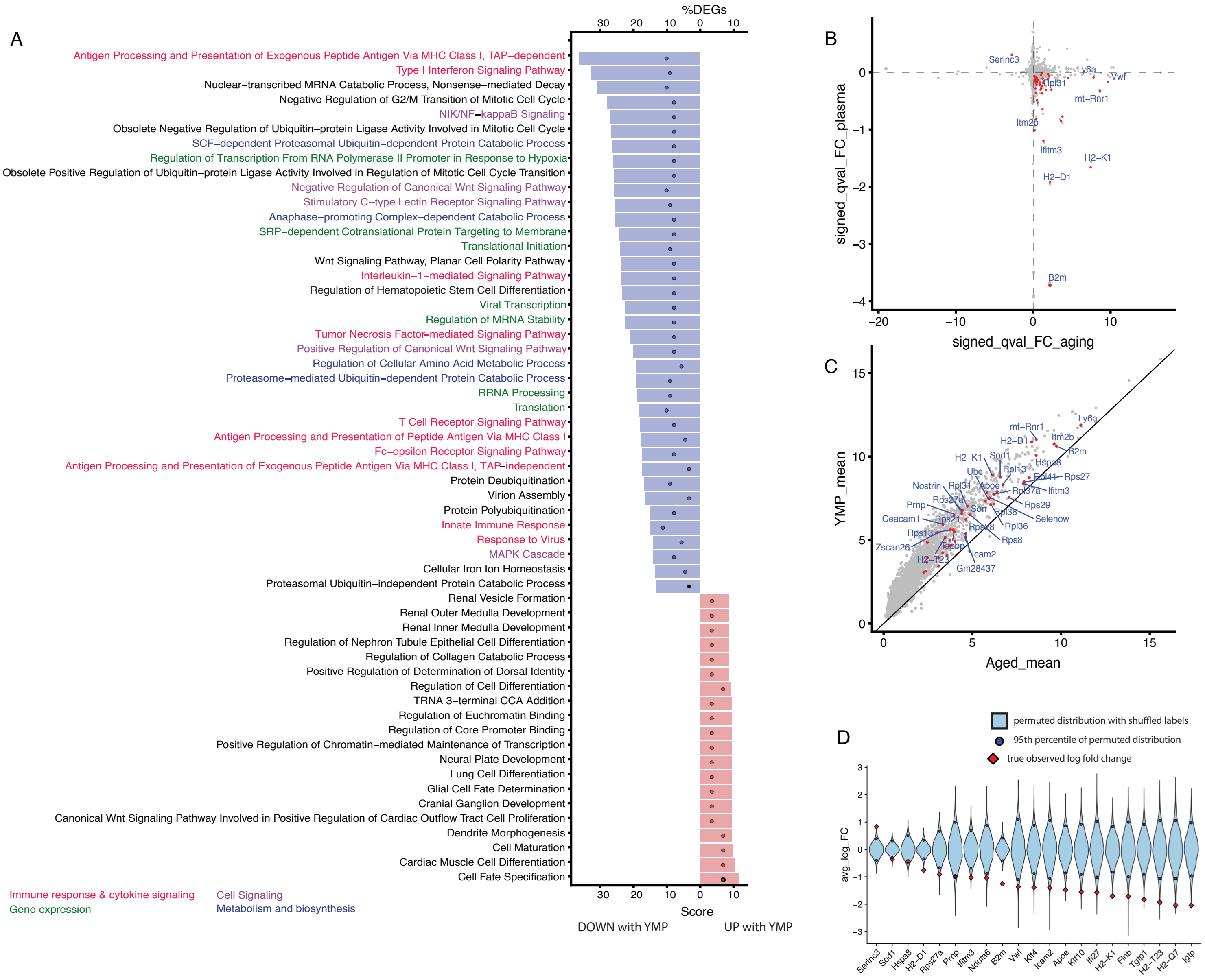


G

SI 9

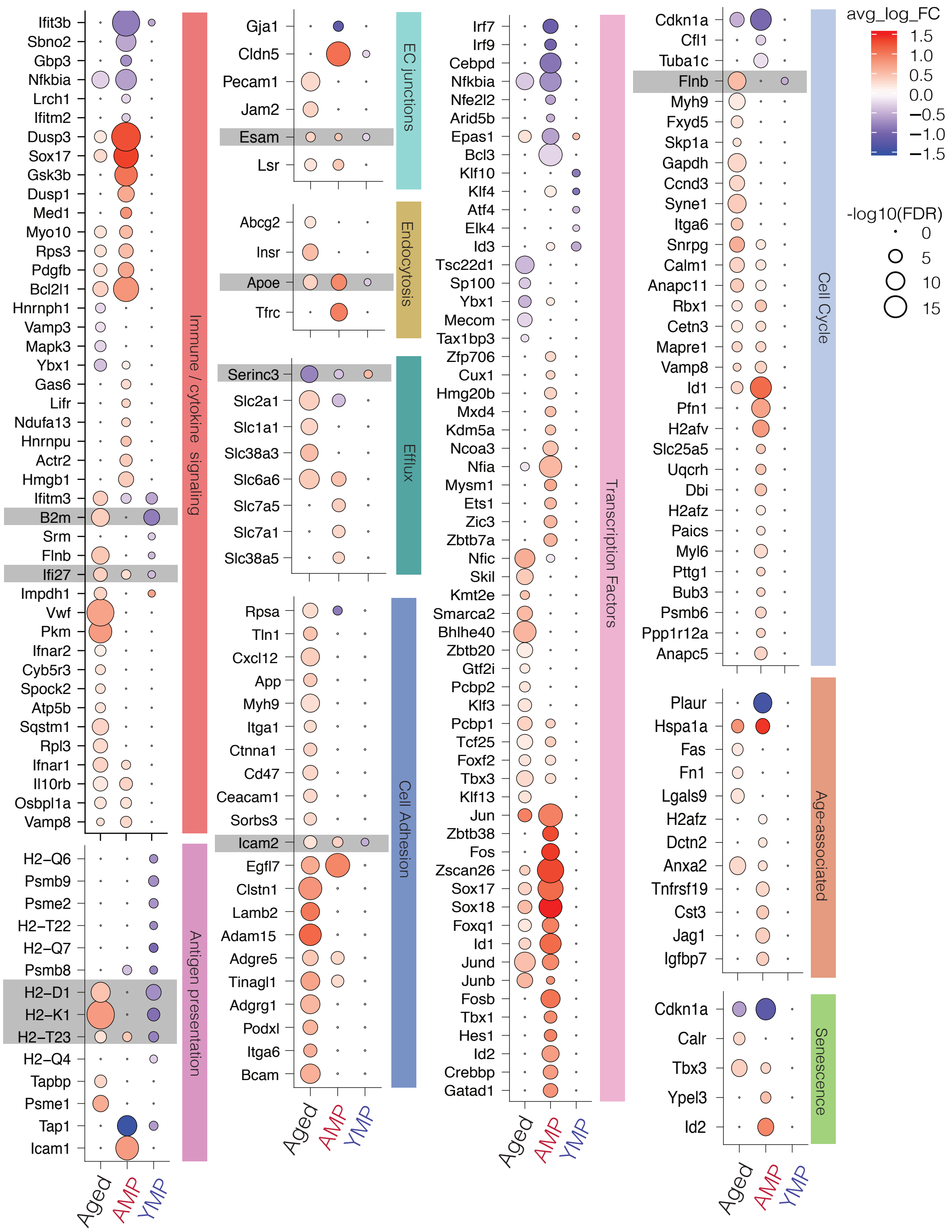
avg_log_FC
1.5
1.0
0.5
0.0
−0.5
−1.0
−1.5
-log10(FDR)
0
5
10
15
Immune / cytokine signaling
Antigen presentation
EC junctions
Endocytosis
Efflux
Cell Adhesion
Transcription Factors
Cell Cycle
Age-associated
Senescence
Aged
AMP
YMP
Ifit3b
Sbno2
Gbp3
Nfkbia
Lrch1
Ifitm2
Dusp3
Sox17
Gsk3b
Dusp1
Med1
Myo10
Rps3
Pdgfb
Bcl2l1
Hnrnph1
Vamp3
Mapk3
Ybx1
Gas6
Lifr
Ndufa13
Hnrnpu
Actr2
Hmgb1
Ifitm3
B2m
Srm
Flnb
Ifi27
Impdh1
Vwf
Pkm
Ifnar2
Cyb5r3
Spock2
Atp5b
Sqstm1
Rpl3
Ifnar1
Il10rb
Osbpl1a
Vamp8
H2−Q6
Psmb9
Psme2
H2−T22
H2−Q7
Psmb8
H2−D1
H2−K1
H2−T23
H2−Q4
Tapbp
Psme1
Tap1
Icam1
Gja1
Cldn5
Pecam1
Jam2
Esam
Lsr
Abcg2
Insr
Apoe
Tfrc
Serinc3
Slc2a1
Slc1a1
Slc38a3
Slc6a6
Slc7a5
Slc7a1
Slc38a5
Rpsa
Tln1
Cxcl12
App
Myh9
Itga1
Ctnna1
Cd47
Ceacam1
Sorbs3
Icam2
Egfl7
Clstn1
Lamb2
Adam15
Adgre5
Tinagl1
Adgrg1
Podxl
Itga6
Bcam
Irf7
Irf9
Cebpd
Nfkbia
Nfe2l2
Arid5b
Epas1
Bcl3
Klf10
Klf4
Atf4
Elk4
Id3
Tsc22d1
Sp100
Ybx1
Mecom
Tax1bp3
Zfp706
Cux1
Hmg20b
Mxd4
Kdm5a
Ncoa3
Nfia
Mysm1
Ets1
Zic3
Zbtb7a
Nfic
Skil
Kmt2e
Smarca2
Bhlhe40
Zbtb20
Gtf2i
Pcbp2
Klf3
Pcbp1
Tcf25
Foxf2
Tbx3
Klf13
Jun
Zbtb38
Fos
Zscan26
Sox17
Sox18
Foxq1
Id1
Jund
Junb
Fosb
Tbx1
Hes1
Id2
Crebbp
Gatad1
Cdkn1a
Cfl1
Tuba1c
Flnb
Myh9
Fxyd5
Skp1a
Gapdh
Ccnd3
Syne1
Itga6
Snrpg
Calm1
Anapc11
Rbx1
Cetn3
Mapre1
Vamp8
Id1
Pfn1
H2afv
Slc25a5
Uqcrh
Dbi
H2afz
Paics
Myl6
Pttg1
Bub3
Psmb6
Ppp1r12a
Anapc5
Plaur
Hspa1a
Fas
Fn1
Lgals9
H2afz
Dctn2
Anxa2
Tnfrsf19
Cst3
Jag1
Igfbp7
Cdkn1a
Calr
Tbx3
Ypel3
Id2
